# Synaptic contributions to cochlear outer hair cell Ca^2+^ homeostasis

**DOI:** 10.1101/2020.08.02.233205

**Authors:** Marcelo J. Moglie, Diego L. Wengier, A. Belén Elgoyhen, Juan D. Goutman

**Affiliations:** Instituto de Investigaciones en Ingeniería Genética y Biología Molecular “Dr. Héctor N. Torres” (INGEBI), C. A. Buenos Aires (1428) Argentina

## Abstract

For normal cochlear function, outer hair cells (OHCs) require a precise regulation of intracellular Ca^2+^ levels. Influx of Ca^2+^ occurs both at the stereocillia tips and through the basolateral membrane. In this latter compartment, two different origins for Ca^2+^ influx have been poorly explored: voltage-gated Ca^2+^ channels (VGCC) at synapses with type II afferent neurons, and α9α10 cholinergic nicotinic receptors at synapses with medio-olivochlear complex (MOC) neurons. Using functional imaging in rodent OHCs, we report that these two Ca^2+^ entry sites are closely positioned, but present different regulation mechanisms. Ca^2+^ spread from MOC synapses is contained by cisternal Ca^2+^-ATPases. Considered a weak drive for transmitter release, we unexpectedly found that VGCC Ca^2+^ signals are comparable in size to those elicited by α9α10 and can be potentiated by ryanodine receptors. Finally, we showed that sorcin, a highly expressed gene product in OHCs with reported Ca^2+^ control function in cardiomy-ocytes, regulates basal Ca^2+^ levels and MOC synaptic activity in OHCs.

## Introduction

Cochlear OHCs are a unique group of cells with a highly polarized structure, featuring a stereocillia bundle on their apical end, and synaptic connections on the basolateral membrane. One important aspect of OHCs physiology is the precise homeostasis and tight regulation of Ca^2+^ during normal activity. Within OHCs stereocilia high concentrations of proteinaceous Ca^2+^ buffers co-exist with large amounts of extrusion pumps, suggesting that mechanisms to quickly clear out Ca^2+^ increments are highly required (Chen et al., 2012; Dumont et al., 2001; Hackney et al., 2005; Sakaguchi et al., 1998; Yamoah et al., 1998). The main Ca^2+^ source are mechanotransducer channels located at the tip of stereocillia (Beurg et al., 2009; Fettiplace & Nam, 2018). A layer of mitochondria right below the cuticular plate (where stereocillia insert) plays the important role of restraining any Ca^2+^ leak into the basolateral compartment of the cell (Beurg et al., 2010; Furness & Hackney, 2006).

Two other important Ca^2+^ sources in OHCs have been less characterized: the voltage gated L-type Ca^2+^ channels (VGCC) (Knirsch et al., 2007), and the α9α10 cholinergic nicotinic receptors (Gómez-Casati et al., 2005; Weisstaub et al., 2002). Both VGCC and α9α10 receptors are located at the basolateral membrane of OHCs, at synapses with type II afferent fibers in the case of the former (Saito, 1990), and on the postsynaptic side of synapses with MOC fibers in the latter (Elgoyhen et al., 1994, 2001). Compared to inner hair cells (IHCs), OHCs show smaller Ca^2+^ currents through VGCC (Beurg et al., 2008; Johnson & Marcotti, 2008; Knirsch et al., 2007; Wong et al., 2013), and also present synaptic ribbons with irregular shapes and fewer vesicles in their vicinity (see for review: Fuchs & Glowatzki, 2015). These contacts have a weak synaptic drive, not suited for sound encoding, but it has been proposed that type II afferents could mediate pain perception (Flores et al., 2015; Liu et al., 2015).

On the other hand, cholinergic MOC synapses onto OHCs are inhibitory and provide the means to modulate mechanosensitivity (Guinan, 1996). Synaptic responses are mediated by the highly Ca^2+^-permeable α9α10 receptors, coupled to the activation of SK2 (Ca^2+^-activated K^+^) channels which ultimately produces inhibition (Fuchs, 1996; Gómez-Casati et al., 2005; Weisstaub et al., 2002). Detailed electron micrographs have shown postsynaptic cisterns within OHCs, closely aligned with presynaptic efferent synaptic contacts (Engström, 1958; Fuchs et al., 2014; Saito, 1980; Smith & Sjöstrand, 1961). This synaptic cistern has been proposed to serve as a Ca^2+^ store that modulates efferent synaptic responses by mechanisms such as Ca^2+^-induced Ca^2+^ release (CICR), through ryanodine receptors (RyR) (Evans et al., 2000; Grant et al., 2006; Lioudyno et al., 2004; Sridhar et al., 1997). Since the evidence for the role of RyR is indirect, in the present study we investigated their participation in directly modulating Ca^2+^ signaling through α9α10 receptors or VGCC. Using an *ex-vivo* preparation of the cochlea from post-hearing onset mice, and functional Ca^2+^ imaging, we found that Ca^2+^ signals from VGCC are unexpectedly large, comparable in amplitude with α9α10 transients, and can be modulated by RyR. On the contrary, Ca^2+^ transients produced by α9α10 activation were not affected by RyR, and were efficiently contained by cisternal Ca^2+^-ATPases.

We also evaluated the role of another regulatory factor, the small Ca^2+^-binding protein sorcin. It has been shown that sorcin controls excitation-contraction coupling in cardiomyocytes, by modulating the activity of RyRs, L-type VGCC and other proteins involved in Ca^2+^ homeostasis (Colotti et al., 2014). More recently, it has been reported that sorcin is among the most differentially expressed genes in OHCs (compared to IHC and supporting cells) (Li et al., 2018; Ranum et al., 2019; Shen et al., 2015), although its role remained hypothetical. By adding sorcin to OHCs cytoplasm produced a strong increase in the resting Ca^2+^ concentration, and inhibition of efferent synaptic currents. Thus, the present results shed light into Ca^2+^ homeostasis in the hair cells involved in sound amplification at the cochlea, and suggest a role for the novel protein sorcin.

## Results

### Local acetylcholine application evokes a global Ca^2+^ rise in OHCs

To directly measure Ca^2+^ influx and spread by activation of α9α10 receptors, the Ca^2+^-sensitive indicator Fluo-4 was loaded into cells through the patch-clamp electrode (Fig. 1A). In a first approach, a local application pipette was used to puff acetylcholine (ACh) onto the organ of Corti preparation. Figure 1B presents a series of images taken at the OHCs base, before and after ACh application (see Fig. 1A for representation of imaging focal plane). The first image in the sequence also shows an array of regions of interest (ROIs) designed to measure fluorescence changes in the cytoplasm as a function of time (%ΔF/F_0_, unless otherwise indicated). The application of a nearly saturating concentration of ACh (1 mM) (Gómez-Casati et al., 2005; Lioudyno et al., 2004) produced a global and long lasting elevation of cytoplasmic Ca^2+^ (images in Fig. 1B and traces of fluorescence intensity as a function of time in Fig. 1C). The mean peak of the ΔF/F_0_ signal was 311 ± 65 %, with a corresponding electrophysiological response integral of 1.9 ± 0.3 nC (n = 6, Fig. 1C-E). These values were considered as an upper limit of efferent activation, since a very high concentration of an externally applied agonist activates receptors distributed throughout the surface of the cell.

**Figure 1.**
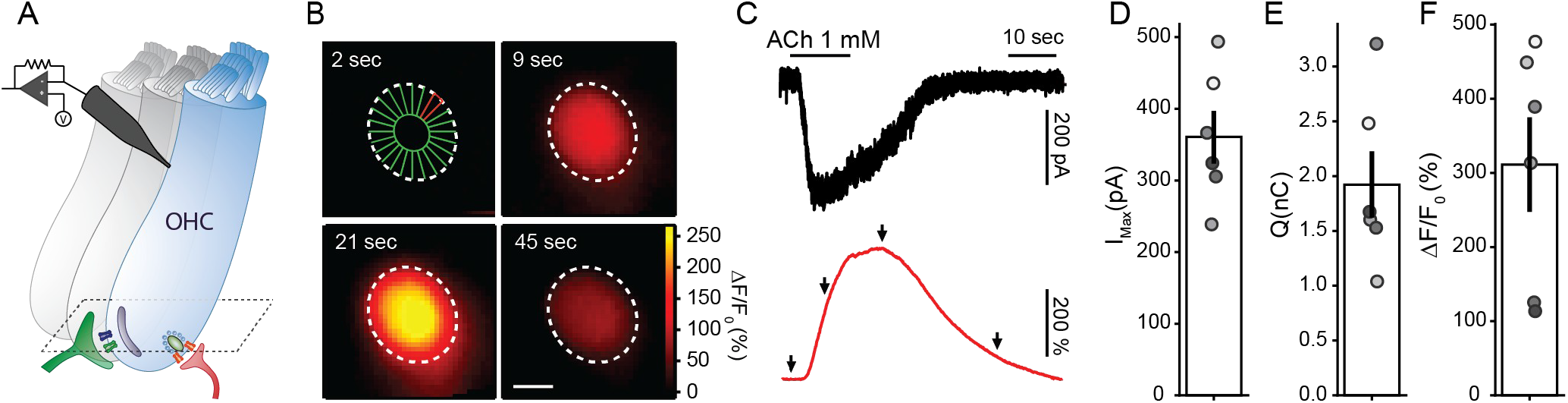
ACh evokes global Ca^2+^ transients in OHCs. A) Illustration of a recorded OHCs within the intact organ of Corti preparation, with synaptic connections (green MOC terminal, red type II afferent), and a scheme of the imaging focal plane (dashed lines). B) Sequence of wide-field microscopy images of an OHC (loaded with Fluo-4) during ACh 1mM perfusion. Dotted white lines represent the outer margin of the OHC’s fluorescence signal. First image shows ROI design scheme in which the cell’s cytoplasm was divided in 24 radial ROIs (see Methods). Scale bar: 5 μm. C) Black trace corresponds to the whole-cell current recorded during ACh perfusion (Vhold =-100mV) and red trace to the ΔF/F_0_ signal measured at the ROI depicted in red on panel B. Arrows indicate the time points of images shown on panel B. D-F) Peak current (D), charge (E), and maximal ΔF/F_0_ (F) during ACh perfusion. Bar plots are mean ± SEM.

### Cholinergic synaptic Ca^2+^ signals in OHCs

An alternative and more physiological approach to investigate efferent input to OHCs was undertaken by electrically stimulating MOC axons innervating these cells. Figure 2A shows a series of images of an OHC during a typical protocol of MOC fibers stimulation. In contrast to that observed with ACh external applications, local and brief Ca^2+^ transients were observed with synaptic activation. Only one Ca^2+^ entry site was observed in each recorded OHC (n = 6 cells). Representative traces of fluorescence changes at the brightest ROI are shown in the bottom panel of Figure 2B (red traces), whereas the corresponding synaptic currents recorded in the same trials are included in the top panel (black traces). α9α10 receptors are highly Ca^2+^ permeable, but due to the much higher concentration of Na^+^ and K^+^ in the extracellular solution (150 mM together vs 1.3 mM Ca^2+^, see Methods), the majority of the current allowed in physiological conditions is due to monovalent cations (Weisstaub et al., 2002). To circumvent the small Ca^2+^ influx, we adopted two technical measures. Firstly, a high affinity Ca^2+^ indicator, such as Fluo-4, was used to detect synaptic Ca^2+^ transients. And secondly, to maximize the Ca^2+^ driving force, OHCs were transiently voltage-clamped at −100 mV in these experiments (for the duration of the synaptic response, otherwise, at −40 mV. See Methods) such that inward currents were due to the activation of both α9α10 receptors and SK2 channels (K^+^ equilibrium potential: ~-82 mV). Paired pulses were used instead of single stimuli to increase the otherwise very low release probability (Ballestero et al., 2011; Vattino et al., 2020). Due to frame rate limitations, the imaging signal appears as the ensemble activation in response to both stimuli in a pair. An average of 100 ± 22 stimulation trials were performed per cell (range 60 – 200, n = 6 cells), with a synaptic success rate of 87 ± 3 % (range: 73 – 98 %, n = 6), judged by the presence of IPSCs (see Methods). In turn, Ca^2+^ signals were detected in 83 ± 5% (range: 64 – 99 %, n = 6) of trials with successful synaptic currents. In paired pulse protocols, an average Ca^2+^ signal of 2.7 ± 0.3 %ΔF/F_0_ (n = 6 cells) was obtained, whereas the integral of the synaptic currents was 7.4 ± 0.8 pC, excluding synaptic failures for the calculation of both averages (n = 6) (Fig. 2C and D). Imaging and electrophysiological responses correlated, as shown in Figure 2E (*r* = 0.88 in this representative example, range: 0.65 – 0.95, n = 6). This strong correlation indicates that the imaging signal is a good proxy for α9α10 and SK2 synaptic activation, most likely reflecting Ca^2+^ influx through nicotinic receptors. Note that failures in imaging events coincided with the smallest IPSCs (black symbols in Fig. 2E).

**Figure 2.**
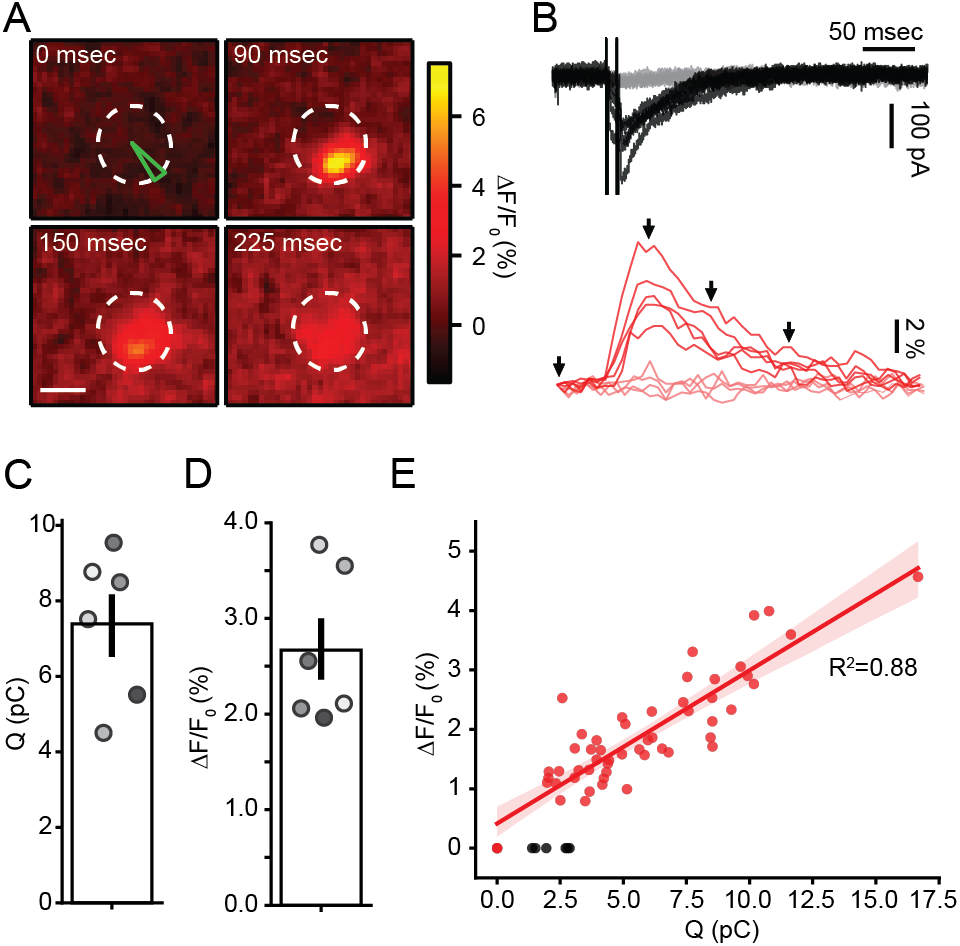
Efferent fiber electrical stimulation evokes localized Ca^2+^ signals in OHCs. A) Sequence of images showing the localized Ca^2+^ increase in an OHC following efferent fiber stimulation. Scale bar: 5 μm. B) *Top*, Representative whole-cell current traces during doble-pulse efferent electrical stimulation at 100 Hz (Vh= −100 mV). Black traces represent those trials where an eIPSC was detected after the stimulus artifact (failures, in gray). *Bottom*, Representative Ca^2+^ transients taken at the ROI indicated in green on panel A, for the same trials shown on the top panel. Red traces correspond to trials were an IPSC was detected and failures in pink. C-D) Mean charge (C) and ΔF/F_0_ (D) for successful efferent fiber stimulation trials. Bar plots are mean ± SEM. E) Size of Ca^2+^ transients as function of charge during efferent stimulation trials in a representative OHC. Black dots represent successful IPSC events with no detectable fluorescence signal.

### Synaptic Ca^2+^ signals during trains of efferent stimuli

MOC neurons neurons can be activated as a result of a feedback loop starting in the afferent pathway and producing steady firing of efferent neurons at rates of up to 200 1/sec (Brown, 1989; Guinan, 2006; Liberman, 1988). Repetitive activation of MOC axons leads to presynaptic facilitation in neurotransmitter release (Ballestero et al., 2011). The postsynaptic consequences of the stimulation in trains are shown as synaptic Ca^2+^ transients in response to repetitive MOC stimulation at 20, 40 and 80 Hz (300 msec of duration each, Fig. 3). Images included in Figure 3A were taken at the peak of the Ca^2+^ rise for each train, whereas panel B shows average synaptic currents (top, in black) and the corresponding Ca^2+^ signals (bottom, in red) taken at the brightest ROI in each cell. Ca^2+^ levels varied with the frequency of the stimulation train, with average amplitudes at 20, 40 and 80 Hz trains of 5.1 ± 1.1 % ΔF/F_0_, 9.5 ± 2.2 % ΔF/F_0_ and 15.6 ± 2.5 % ΔF/F_0_ (n = 8, p = 0.0001 Friedman’s test, Fig. 3C), respectively. The integral of the ensemble synaptic response across the duration of the train was 23.6 ± 6.2 pC, 43.3 ± 11.0 pC and 64.1 ± 8.8 pC (n = 8, p < 0.0001 Friedman’s test, Fig. 3D).

**Figure 3.**
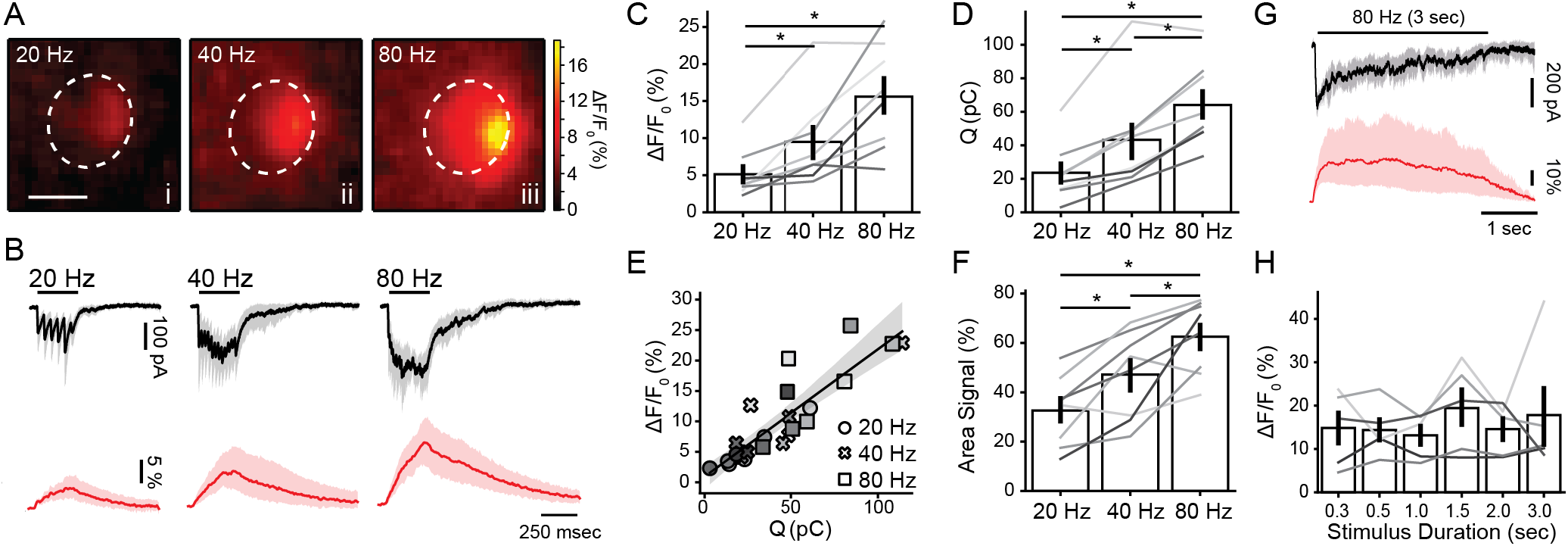
Amplitude and spread of efferent Ca^2+^ transients are dependent upon stimulation frequency. A) Representative images of an OHC at the peak of the fluorescence signal during efferent fiber electrical stimulation at 20 (i), 40 (ii) and 80 (iii) Hz. B) Mean inhibitory current traces (black) recorded during 300 msec efferent fiber electrical stimulation at 20, 40 and 80 Hz (Vh= −100 mV). Red traces show the mean ΔF/F_0_ at the ROI with the highest fluorescence signal. Peak Ca^2+^ values (C) and charge (D) for 300 msec stimulation trains at tested frequencies. E) Amplitude of fluorescence signal as a function of charge. Each symbol represent a different stimulation frequency. F) Spread of the Ca^2+^ signal within each OHC cytoplasmic space (as percentage of area in the imaged plane). Values were taken at the time point where signal peaked. G) Mean whole-cell synaptic response (black) and Ca^2+^ signals (red) obtained during 3 sec electrical stimulation of efferent fibers at 80 Hz. H) Peak fluorescence amplitude for trains with duration between 0.3 and 3 seconds (at 80Hz). Bar plots are mean ± SEM. Friedman’s Test, * p<0.05.

A close interdependence of synaptic and Ca^2+^ responses was also noted in the correlation between peak ΔF/F_0_ and the integral of synaptic currents, elicited by trains at different frequencies (*r* = 0.88, Fig. 3E). As suggested by the representative images, trains at 20 Hz elicited a localized Ca^2+^ rise with a measurable spread which accounted for 31 ± 5 % of the area corresponding to the imaged OHC area. At 40 or 80 Hz Ca^2+^ signals reached a larger part of the cytoplasmic space, with averages values of 48 ± 6 % and 63 ± 5 %, respectively (n = 8, p < 0.0001 Friedman’s test, Fig. 3F).

In order to obtain a better understanding of Ca^2+^ dynamics during sustained MOC activity, trains of stimuli at 80 Hz were applied for longer periods of time, up to 3 sec (Fig. 3G and H). A sustained Ca^2+^ load was observed in OHCs, with peak values that did not differ when different train durations were compared (p = 0.57 Friedman’s test). This latter result indicates that mechanisms for controlling Ca^2+^ entering from efferent sources are highly efficient, preventing a large Ca^2+^ load even during a 3 sec stimulation. The role of the sub-synaptic cistern in this phenomenon has been suggested in the past (Evans et al., 2000; Lioudyno et al., 2004; Sridhar et al., 1997) and it was evaluated in the following section.

Comparing Ca^2+^ signals in Figs. 1 and 3 it becomes clear that synaptic activation of α9α10 receptors during trains is (at least) ~20 times smaller than that obtained with locally applied ACh. This could be due to the rather low release probability of MOC terminals (Ballestero et al., 2011; Vattino et al., 2020), to the fact that the stimulation paradigm typically activates a single axon (see Methods), and that activation by ACh does not rely on vesicle availability. This drastic difference in measurable ΔF/F_0_ amplitude between ACh application and electrical stimulation also suggests that the Ca^2+^ indicator used in these experiments, Fluo-4, does not saturate during synaptic trains. The tight correlation between ΔF/F and charge in Fig. 3E further supports this conclusion.

### Modulation of efferent Ca^2+^ by cisterns

One important factor are sarcoplasmic/endoplasmic reticulum Ca^2+^-ATPases (SERCA), responsible for removing free Ca^2+^ ions from the cytoplasm. To address the role of SERCA in shaping Ca^2+^ transients, the specific blocker thapsigargin (1 μM) was added to the bath during an OHC recording. In the absence of any stimulation, the basal fluorescence increased when perfusing thapsigargin from a control value of 1396 ± 551 A.U. (arbitrary units) to 1851 ± 502 A. U. (n = 6, p = 0.024 Wilcoxon test) (Fig. 4B inset). This ~ 30% increase in the basal cytoplasmic concentration of Ca^2+^ would represent a basal SERCA activity responsible for pumping ions out of the OHC cytoplasm at rest.

**Figure 4.**
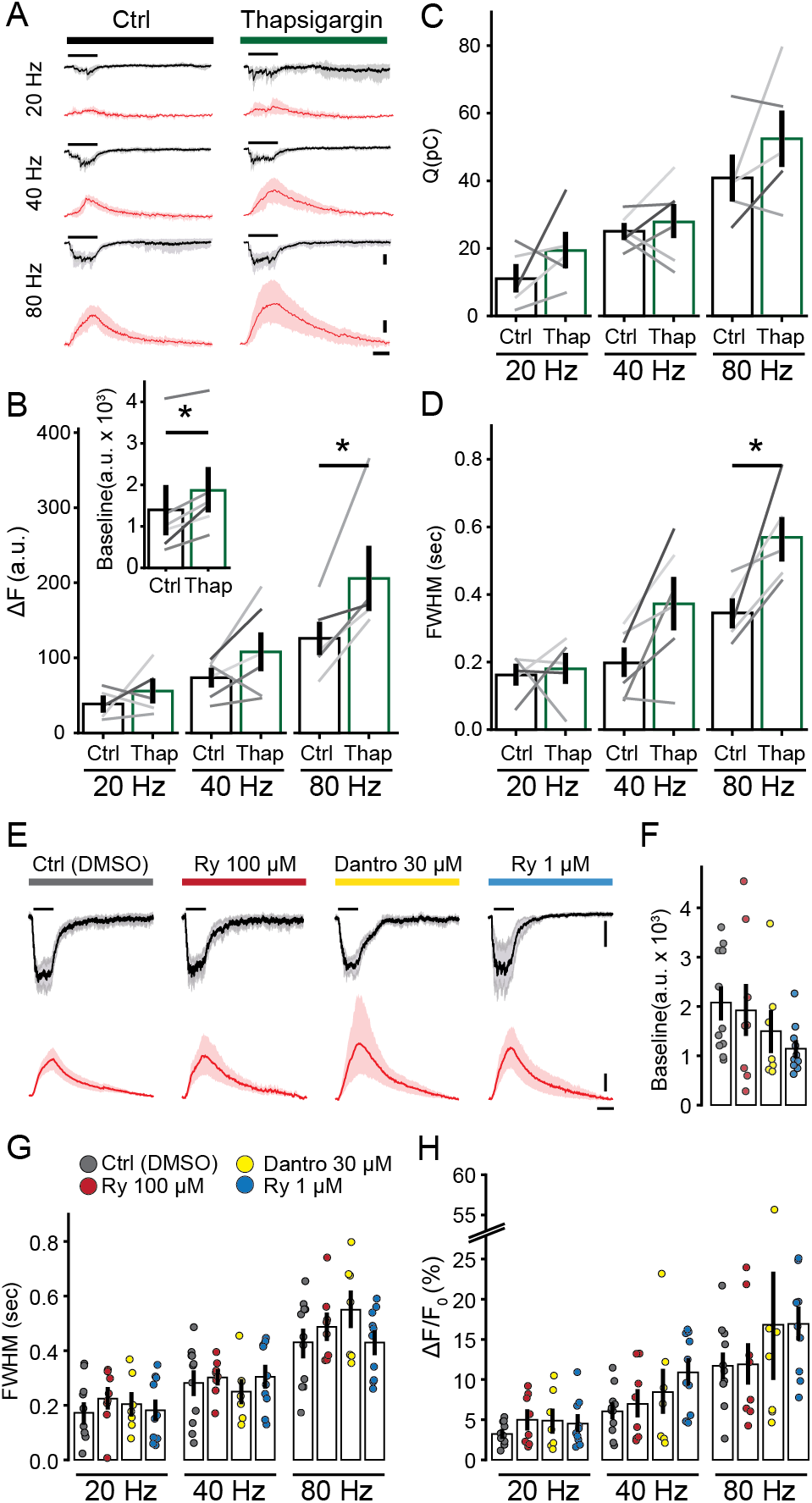
Efferent Ca^2+^ signals are modulated by cisternal ATPases, but not ryanodine receptors. A) Mean synaptic responses (black) and Ca^2+^ transients (red) during 300 msec electrical stimulation at 20,40 and 80 Hz before and after perfusion of Thapsigargin. B) Peak of Ca^2+^ transients (as ΔF), C) Charge of synaptic responses and D) duration of Ca^2+^ transients (as full width at half maximum, FWHM). Inset: Baseline fluorescence signal before and after perfusion of Thapsigargin. E) Synaptic currents (mean, black) and Ca^2+^ transients (red) obtained during efferent fibers stimulation (300 msec, 80 Hz), using an intracellular solution containing vehicle (DMSO), RyR blockers (Ryanodine 100 μM and Dantrolene 30 μM) and a RyR agonist (Ryanodine 1 μM). F) Baseline fluorescence for each condition. G) Duration (FWHM) and H) maximal fluorescence signal (as ΔF/F_0_) for each intracellular solution. Bar plots are mean ± SEM. Wilcoxon signed-rank test, * p<0.05.

To determine the role for SERCA in regulating synaptic Ca^2+^, trains of MOC stimuli were applied before and during the application of thapsigargin (Fig. 4A-D). Peak ΔF values were calculated (Fig. 4B), instead of ΔF/F_0_, to avoid the artifact effect of a higher basal fluorescence on the estimation of the size of the Ca^2+^ transients. Synaptic currents integral are shown in Figure 4C. Peak ΔF values for control trains at 20, 40 and 80 Hz were respectively 38.4 ± 8.6 A. U., 73.4 ± 10.5 A. U., and 125.9 ± 21.8 A. U. (n = 5), showing statistical differences between trains as previously indicated for ΔF/F_0_ measure (p = 0.0085, Friedman’s test). With thapsigargin in the bath, ΔF values grew to 55.8 ± 14.1 A. U., 107.9 ± 24.5 A. U., and 205.7 ± 39.9 A. U. respectively for the 20, 40 and 80 Hz trains (n = 6), showing statistically significant differences for the 80 Hz train compared to control (p = 0.04, Wilcoxon signed-rank test).

The effect of thapsigargin on the duration of the synaptic Ca^2+^ transients was also analyzed, by measuring the ‘full width at half maximum’ (FWHM) of the signal. In control conditions, FWHM values varied with the stimulation frequency: 211 ± 45 ms at 20 Hz, 344 ± 80 ms at 40 Hz, and 485 ± 79 ms at 80 Hz (n = 5, p = 0.0097, Friedman’s test). In the case of the 20 Hz train, the FWHM of the transient was even shorter than the stimulating train duration (300 ms) due to sporadic synaptic activation. Thapsigargin prolonged Ca^2+^ transients with average values of 569 ± 62 ms at 80 Hz, 372 ± 39 ms at 40 Hz, and 180 ± 20 ms at 20 Hz (n = 5, p = 0.048 Wilcoxon signed-ranked test for the 80 Hz train compared to control). Taken together, these results indicate that SERCA pumps, and the sub-synaptic cistern, play a significant role in accelerating the removal of Ca^2+^ entering through efferent synapses, and also in curtailing the peak of the transient.

Since the presence of RyR has been shown both morphologically and functionally at the MOC - OHC synapse (Evans et al., 2000; Grant et al., 2006; Lioudyno et al., 2004), in the following experiments we tested Ca^2+^ dynamics in the presence of drugs that modulate RyR. To avoid indirect or presynaptic effects, drugs were included in the patch pipette (control experiments included vehicle – DMSO). Low (1 μM) and high (100 μM) concentrations of ryanodine were used to either activate or block RyR. Dantrolene, a specific inhibitor of these receptors, was also used. Interestingly, none of these treatments showed any effect on either the amplitude of synaptic Ca^2+^ transients (Fig. 4E and H, Kruskal-Wallis test) (same for ΔF), duration (estimated as FWHM, Fig. 4E and G, Kruskal-Wallis test) or basal Ca^2+^ (Fig. 4F, Kruskal-Wallis test). The corresponding values for these experiments are shown in Table 1.

**Table I.** Values for integral of synaptic currents, amplitude and duration Ca^2+^ signals during step depolarization of OHCs or MOC stimulation at 20, 40 and 80 Hz trains, in different pharmacological conditions.

|  |  | **Q (pC)** | **$\Delta F/F_0$ (%)** | **FWHM (sec)** |
| --- | --- | --- | --- | --- |
| **Depolarization** | Ctrl (DMSO) | - | $5 \pm 1$ (n=11) | $0.23 \pm 0.04$ (n=8) |
| | Ryanodine 100 μM | - | $13 \pm 6$ (n=8) | $0.22 \pm 0.07$ (n=8) |
| | Dantrolene 30 μM | - | $13 \pm 5$ (n=7) | $0.27 \pm 0.07$ (n=7) |
| | Ryanodine 1 μM | - | $16 \pm 3$ (n=10) | $0.35 \pm 0.02$ (n=10) |
| **20 Hz** | Ctrl (DMSO) | $20 \pm 4$ (n=11) | $3 \pm 0$ (n=11) | $0.17 \pm 0.03$ (n=11) |
| | Ryanodine 100 μM | $21 \pm 4$ (n=8) | $5 \pm 1$ (n=8) | $0.22 \pm 0.04$ (n=8) |
| | Dantrolene 30 μM | $13 \pm 2$ (n=7) | $5 \pm 1$ (n=7) | $0.20 \pm 0.04$ (n=7) |
| | Ryanodine 1 μM | $17 \pm 5$ (n=10) | $5 \pm 1$ (n=10) | $0.18 \pm 0.04$ (n=10) |
| **40 Hz** | Ctrl (DMSO) | $34 \pm 5$ (n=11) | $6 \pm 1$ (n=11) | $0.28 \pm 0.04$ (n=11) |
| | Ryanodine 100 μM | $32 \pm 4$ (n=8) | $7 \pm 2$ (n=8) | $0.30 \pm 0.02$ (n=8) |
| | Dantrolene 30 μM | $27 \pm 3$ (n=7) | $8 \pm 3$ (n=7) | $0.25 \pm 0.04$ (n=7) |
| | Ryanodine 1 μM | $37 \pm 6$ (n=10) | $11 \pm 2$ (n=10) | $0.30 \pm 0.04$ (n=10) |
| **80 Hz** | Ctrl (DMSO) | $61 \pm 6$ (n=11) | $12 \pm 2$ (n=11) | $0.43 \pm 0.05$ (n=11) |
| | Ryanodine 100 μM | $48 \pm 6$ (n=8) | $12 \pm 3$ (n=8) | $0.49 \pm 0.04$ (n=8) |
| | Dantrolene 30 μM | $46 \pm 4$ (n=7) | $17 \pm 7$ (n=7) | $0.55 \pm 0.07$ (n=7) |
| | Ryanodine 1 μM | $60 \pm 7$ (n=10) | $17 \pm 2$ (n=10) | $0.43 \pm 0.04$ (n=10) |

### Efferent Ca^2+^ regulation by sorcin

A number of genes related to Ca^2+^ regulation are highly-expressed in OHCs (compared to IHCs and supporting cells) (Li et al., 2018; Ranum et al., 2019; Shen et al., 2015). Sorcin, a gene product related to the regulation of CICR in cardiac myocytes (Farrell et al., 2003), was identified in OHCs both by RNAseq and inmunostaining (Li et al., 2018; Ranum et al., 2019; Shen et al., 2015). We evaluated the effect of sorcin on efferent Ca^2+^ transients, by adding recombinant sorcin (3 μM) to the intracellular pipette solution. The basal Ca^2+^ concentration in the OHC cytoplasm was higher in the presence of sorcin. In control conditions, basal fluorescence was 1145 ± 112 A. U., whereas with sorcin this value increased to 2091 ± 316 A. U (Fig. 5B, n = 6, p = 0.02 Mann-Whitney U test). Considering that basal fluorescence was partially dependent on the recording conditions of a given cell, and that sorcin recordings were done in a different set of cells than controls, the possibility that cells with sorcin were more leaky than the controls was evaluated. Basal fluorescence as a function of leak current (within the first 10 minutes of recording) was plotted and grouped in control and sorcin cells (Fig. 5C). Each group was fitted with a line showing that basal fluorescence grew faster in sorcin cells and with a significantly larger slope (F-test, p = 0.0038) within a similar range of leak current values, implicating that sorcin increased resting Ca^2+^ concentration values *per se*, and not due to worsened recording conditions.

**Figure 5.**
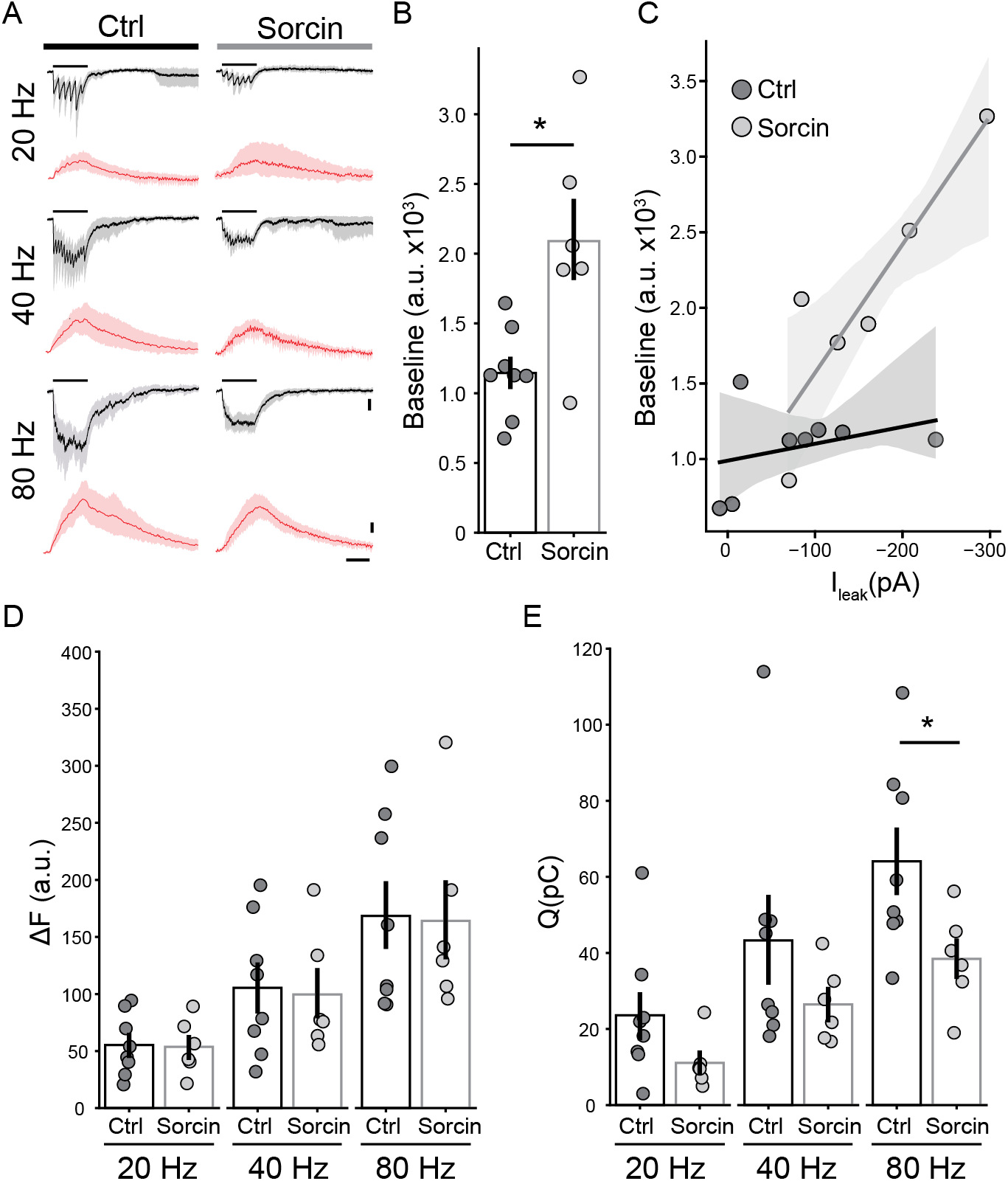
Sorcin produced a rise in resting Ca^2+^ levels and inhibited efferent synaptic currents. A) Mean traces of ensemble synaptic currents (black) and Ca^2+^ transients (red) in OHCs obtained by 300 msec electrical stimulation of efferent fibers at 20,40 and 80 Hz in control conditions (Ctrl) and with the addition of sorcin to the intracellular solution. Scale bars: 50 pA, 25 A. U. and 200 msec. B) Baseline fluorescence in control conditions and in the presence of sorcin. C) Baseline fluorescence as a function of leak current (I_leak_) for each recorded cell during the first 10 minutes of recording. Linear fits were performed separately for control and sorcin groups. D) Peak of Ca^2+^ signals (ΔF) for different train frequencies and intracellular conditions. E) Integral of synaptic responses (Q) during trains of stimuli for each condition. Bar plots are mean ± SEM. Mann-Whitney test, * p<0.05.

Responses to electrical stimulation in sorcin experiments (Fig. 5) were represented in ΔF, instead of ΔF/F_0_, to avoid artifact effects due the higher basal fluorescence, as previously indicated. As shown in Figure 5D, the mean amplitude of Ca^2+^ transients evoked with different stimulation frequencies did not differ in the presence of sorcin (20 Hz trains: control = 55 ± 10 A. U. (n = 8) – sorcin = 54 ± 10 A. U. (n = 6); 40 Hz: control = 105 ± 21 A. U. (n = 8) – sorcin = 100 ± 21 A. U. (n = 6); 80 Hz: control = 169 ± 30 A. U. (n = 8) – sorcin = 164 ± 34 A. U. (n = 6) Mann-Whitney U test). In the presence of sorcin FWHM did not differ from the control (not shown). Interestingly, and despite unchanged Ca^2+^ signals, a strong reduction in synaptic currents of up to ~50% was observed with sorcin (Fig. 5E). The integral of these synaptic responses was calculated, obtaining average values with sorcin of 11.1 ± 2.8 pC for the 20 Hz trains, 26.5 ± 4.0 pC for 40 Hz, and 38.4 ± 5.1 pC for 80 Hz trains (control values already mentioned above) (n = 6, p = 0.0293, Mann-Whitney U test for the 80 Hz trains). The reduction of synaptic currents produced by sorcin is most likely due to a reduction in the SK2 component of the response, since: i) at −100 mV both the nicotinic and SK2 components are inward, and ii) α9α10 receptors response seems to be unchanged, according to ΔF values in Figure 5D that are entirely due to the activation of the nicotinic receptors. Thus, even without affecting Ca^2+^ influx, sorcin produced a reduction in the efferent inhibitory action.

### Ca^2+^ entry through L-type Ca^2+^ channels

Ca^2+^ influx through VGCC was investigated applying step depolarizations to OHCs, from a holding potential of −100 mV, up to a range of potentials between −30 and +30 mV. Representative Ca^2+^ transients are shown in Figure 6A, with maximum values that followed the expected bell-shaped dependence on membrane potential, peaking at +10/+20 mV (Fig. 6B).

**Figure 6.**
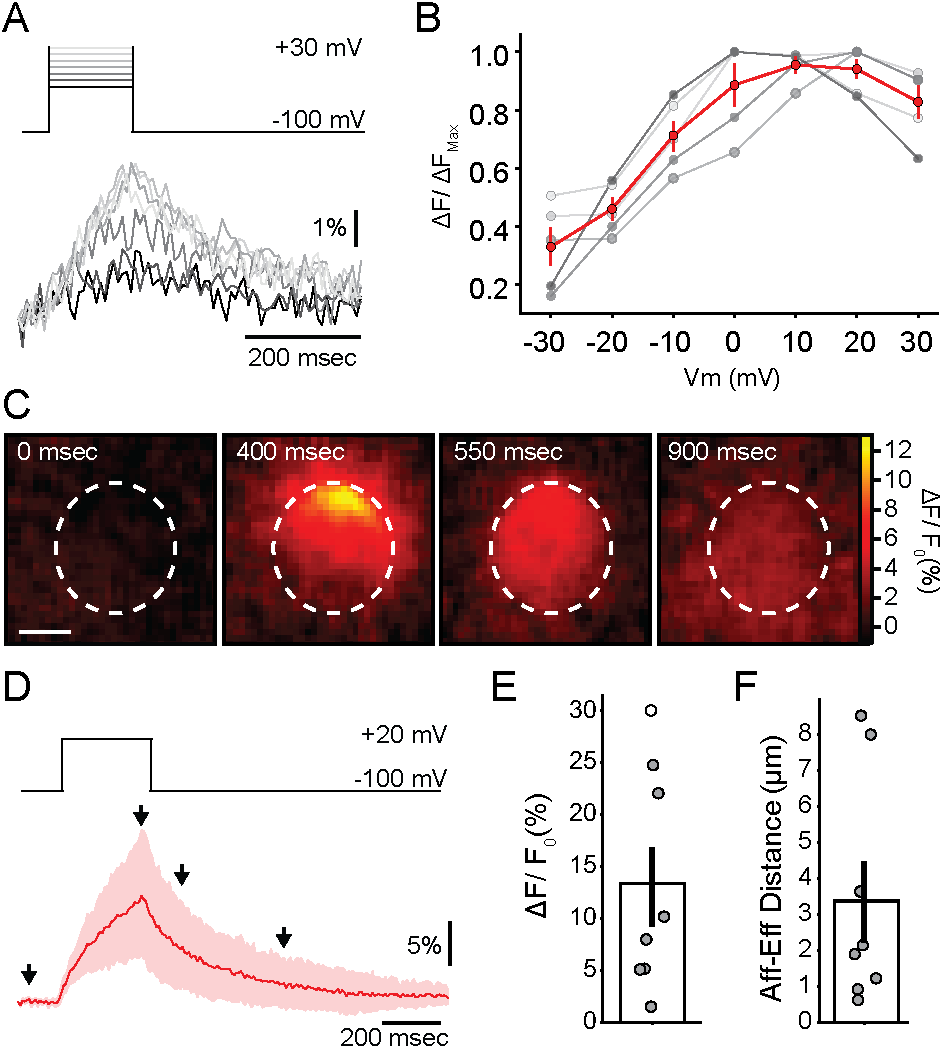
Ca^2+^ transients during VGCC activation in OHCs. A) Representative traces of Ca^2+^ transients measured at the brightest ROI, for voltage steps between −30 and + 30 mV. B) Normalized maximum for Ca^2+^ transients in each cell (gray symbols) as a function of voltage. Red trace and marks represent mean ± SEM. C) Sequence of images showing the localized Ca^2+^ concentration increase in an OHC during 300 msec depolarization to +20 mV. Scale bar: 5 μm. D) Mean trace of the Ca^2+^ signal (in red) during a step pulse to +20 mV (top panel). E) Average ΔF/F_0_ for +20 mV steps in OHCs. F) Average distance between locations of afferent (VGCC) and efferent (MOC) Ca^2+^ signals within each OHC. Bar plots are mean ± SEM. Friedman’s Test, * p<0.05.

Step depolarizations to +20 mV for 300 msec (same duration of MOC train stimulation in Figs. 3–5) evoked Ca^2+^ transients with an average peak of 13.3 ± 3.8 % ΔF/F_0_, that unexpectedly matched in size with those obtained with 40 and 80 Hz trains of MOC stimulation (see Figs. 3C and 6C-E) (n = 8, p > 0.05 Friedman’s test). Moreover, the spread of the fluorescence signal also resembled that observed with 40 and 80 Hz efferent trains (57 ± 7 % of OHC area, p > 0.05 Friedman’s test).

Previous evidence indicates that type II afferents on OHCs are closely positioned with efferent synapses and sub-synaptic cisterns (Fuchs et al., 2014; Saito, 1980, 1990). A functional quantification of this proximity was evaluated by measuring the distance between locations of VGCC and MOC Ca^2+^ transients within a given cell (see Methods) (Fig. 6C and F). An average value of 3.7 ± 1.1 μm (n = 8) was estimated, ranging from 0.6 to 8.5 μm, with 5/8 cells with values ≤ 2 μm. The possibility of modulation of VGCC Ca^2+^ signals by cisterns is further investigated in the following section.

### Modulation of Ca^2+^ influx through VGCC by ryanodine and sorcin

Thapsigargin, ryanodine and dantrolene were used to evaluate the role of SERCA pumps and RyR in modulating depolarization evoked Ca^2+^ transients. Thapsigargin application in the bath did not produce any change in the amplitude of the depolarization evoked Ca^2+^ signal (Figs. 7A and B) (control ΔF: 68 ± 24 A. U., thapsigargin ΔF: 53 ± 9, n = 5, p = 0.58 Wilcoxon signed-rank test). At 1 μM ryanodine, a concentration that activates RyR, a strong potentiation of the Ca^2+^ transient was observed, with average peak values of 15.8 ± 3.5 %ΔF/F_0_, compared to control (vehicle) 4.6 ± 1.6 %ΔF/F_0_ (n = 10, p < 0.05 Kruskal-Wallis test) (Figs. 7C and D). Neither 100 μM ryanodine nor dantrolene showed any modulatory effect (12.6 ± 6.5 % ΔF/F_0_ (n = 8), and 13.2 ± 4.7 % (n = 7), respectively; p > 0.05 Kruskal-Wallis test).

**Figure 7.**
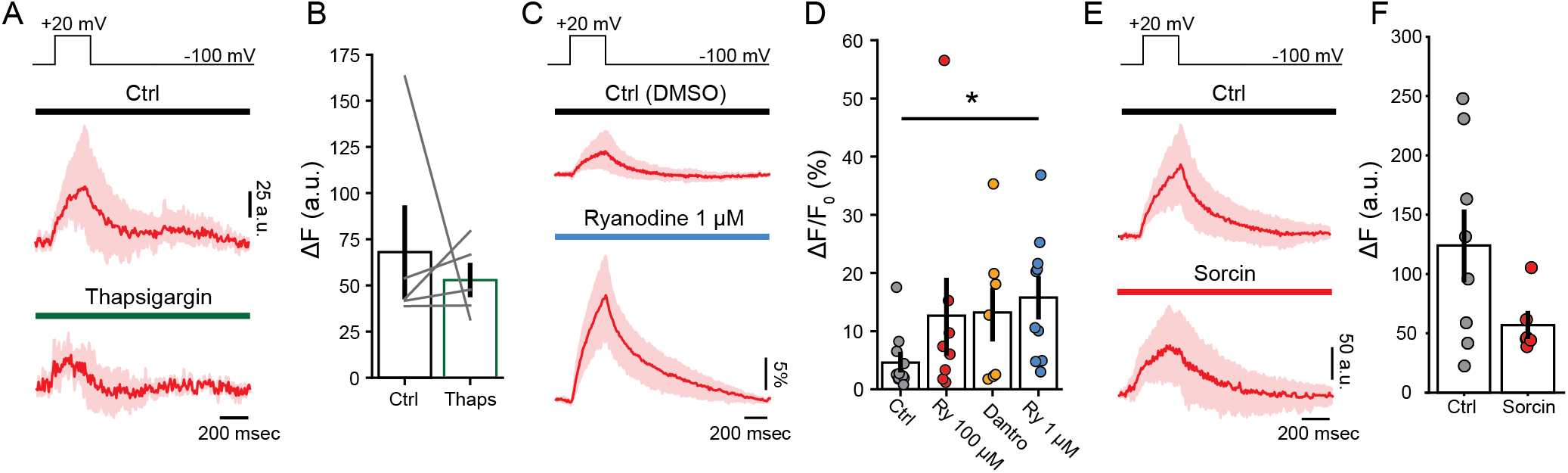
Modulation of afferent Ca^2+^ influx through VGCC by ryanodine and sorcin. A) Mean traces of the Ca^2+^ transients obtained with 300 msec depolarization to +20 mV before (top) and during perfusion of Thapsigargin (bottom). B) Average peak Ca^2+^ level (ΔF) for step depolarizations as in A). C) Mean traces of the Ca^2+^ transients during with steps to +20 mV, using intracellular solutions containing DMSO or Ryanodine 1 μM. D) Average peak Ca^2+^ signal (ΔF/F_0_) for experiments in C), and also with an antagonistic concentration of ryanodine (Ry) (100 μM) and dantrolene (30 μM). Wilcoxon signed-rank test, * p<0.05. E) Mean ΔF traces obtained in OHCs loaded with sorcin protein or control. F) Mean peak fluorescence signal (ΔF) for experimens in E). Bars are mean ± SEM.

Finally, the effect of sorcin on Ca^2+^ transients produced by VGCC is shown in Figure 7E and F. A strong tendency to a reduction in peak ΔF values was observed (control: 124.0 ± 30.0 A. U., sorcin: 57.0 ± 10.2 A. U. n = 6). However, due to the high variability observed in control experiments (range: 22-200 A. U.) this difference did not reach statistical significance (p = 0.228, Mann-Whitney U test).

## Discussion

The present results provide a direct analysis of synaptic Ca^2+^ dynamics in OHCs during both MOC (efferent) and VGCC (afferent) activation. It also shows evidence for cisternal modulation of amplitude, spread and duration of Ca^2+^ transients. In addition, this study demonstrates for the first time a functional role for sorcin in OHCs, a Ca^2+^ binding protein with a well described role in regulating Ca^2+^ concentration in cardiomyocytes cytoplasm (Farrell et al., 2003).

The amplitude of Ca^2+^ transients shows a strong dependence on MOC stimulation at frequencies between 20 and 80 Hz (Fig. 3), which might explain the reported changes in MOC inhibitory strength as a function of stimulation rate (Art et al., 1984; Galambos, 1956; Gifford & Guinan, 1987). Significantly, only one Ca^2+^ hotspot was found per OHC, suggesting that a single MOC fiber is stimulated at a time, although more are present (Liberman, 1990; Warr, 1992). A single Ca^2+^ spot was also observed when activating VGCC by step depolarizations, although multiple contacts with type II afferent coexist in one OHC within a close proximity (Fuchs & Glowatzki, 2015). How afferent synapses in OHCs are activated *in vivo* is still not known, but our experiments show that Ca^2+^ signals produced by VGCC are unexpectedly high, similar in amplitude to those through α9α10, and can be further potentiated by RyR action.

Synaptic Ca^2+^ signals of both origins were largely smaller than those obtained by local perfusion of a high ACh concentration (compare Figs. 1, 3 and 6), suggesting that the Ca^2+^ indicator used in these experiments (Fluo-4) was not saturated during physiological activation. The selection of the indicator was based upon the signal to noise ratio obtained in experiments that sought to detect Ca^2+^ influx through the highly Ca^2+^ permeable α9α10 receptors, which actually allows predominantly Na^+^ through when activated in physiological conditions (140 mM Na^+^ and 1.3 mM Ca^2+^, as in our experiments) (Moglie et al., 2018; Weisstaub et al., 2002).

### Ca^2+^ homeostasis in OHCs and efferent regulation

Several studies report alternative mechanisms that operate in OHCs to handle Ca^2+^ (Fettiplace & Nam, 2018). Millimolar concentrations of two different buffers are present in the cytoplasm of OHCs: oncomodulin (also known as parvalbumin-β) and calbindin-D28k, with the highest values observed at the base of the cells, where synaptic contacts reside (Hackney et al., 2005; Sakaguchi et al., 1998). In the oncomodulin knock-out mouse, a progressive OHCs loss is observed that leads to cochlear dysfunction (Tong et al., 2016). A similar phenomenon is observed in the knock-out of PMCA2 Ca^2+^-ATPases (Giacomello et al., 2011), that are responsible for pumping out Ca^2+^ ions that enter through MET channels even in quiet (Johnson et al., 2011; Tucker & Fettiplace, 1995; Yamoah et al., 1998). These results highlight the importance of Ca^2+^ handling in maintaining the integrity of OHCs and cochlear function.

The control of Ca^2+^ diffusion is further exerted by mechanisms such as Na^+^-Ca^2+^ exchangers and mitochondrial uptake, although they both operate on a time scale of hundreds of milliseconds, slower than buffers and PMCA2 pumps (Beurg et al., 2010; Ikeda et al., 1992; Nicholls, 2005). In OHCs, mitochondria are abundantly found in a layer right below the cuticular plate, playing the important role of containing Ca^2+^ leak into the basolateral compartment of the cell (Beurg et al., 2010; Furness & Hackney, 2006). Similarly, on the basal pole of OHCs, the large cistern located in physical opposition to MOC vesicle releasing sites, would not only operate as a barrier to prevent free diffusion of Ca^2+^ entering through nicotinic receptors, but has the additional role of a ‘Ca^2+^ sponge’ that removes free ions out the cytoplasm and shorten synaptic responses (Fig. 4). Taking into account the tight functional coupling between α9α10 and SK2 (Oliver et al., 2000) and the small cytoplasmic space between plasma and cisternal membranes (Fuchs et al., 2014), a relatively small Ca^2+^ influx would effectively produce synaptic inhibition through SK2 activation. However, since efferent fibers operate best at prolonged MOC activation (Brown, 1989; Robertson & Gummer, 1985), one could propose that cisterns function to control excessive Ca^2+^ spread and prevent its spill over. Evidence from Figure 4, using train stimulation to MOC fibers, suggests that cisterns are responsible not only for speeding up the decay of efferent Ca^2+^ but also limiting its spread. However, taking into account the volume of this small domain and the spread of the Ca^2+^ transients (Fig. 3F), it is very likely that Ca^2+^ signals evoked at high frequency MOC stimulation result, at least partially, from Ca^2+^ that escaped into the cytoplasm. This might differ in conditions with more intact intracellular buffers concentrations. Due to the fragility of the tissue, as well as the instability of OHCs recordings from older animals, we decided to circumscribe our experiments within the P12-P14 age range. Therefore, some of our conclusions may differ in more mature OHCs.

Our results do not provide support for RyR participation during MOC activation within a rather wide range of stimulation strengths, 20 to 80 Hz trains of 300 msec of duration. This result contrasts with previous observations in OHCs in which the role of RyR was evaluated, not directly on Ca2+ transients as we did here, but by measuring efferent synaptic activation (Lioudyno et al., 2004). It is possible though, that RyR are engaged during MOC activation only when a strong preceding hair cell excitation occurred, as suggested previously (Im et al., 2014; Zachary et al., 2018). Whether this phenomenon occurs *in vivo* remains to be proven, but is supported by the observation that MOC inhibition is potentiated by a preceding auditory stimulation, that could produce the required excitation to hair cells (Kujawa & Liberman, 1999).

### Ca^2+^ influx through VGCC and type II afferent activation

Ca^2+^ currents through VGCC in OHCs are several fold smaller than in IHCs and present a shift to the right in the current-voltage relation (Johnson & Marcotti, 2008; Knirsch et al., 2007; Wong et al., 2013) Accordingly, Ca^2+^ signals in Figure 6 peaked at +20 mV (−20 mV for IHCs), and their amplitudes are three-fold smaller (13.3 % ΔF/F_0_, with a 300 ms step) compared to brief depolarizations (20 msec) to sub-maximal potentials (−30 mV) in IHCs (42 % ΔF/F_0_) (Moglie et al., 2018). Other features of afferent synapses in OHCs further indicate that the synaptic drive is small in these cells when compared to IHCs. Ribbons in active zone areas with type II afferents are small, irregular in shape, and are surrounded by few vesicles (Fuchs & Glowatzki, 2015). However, our results show that Ca^2+^ transients elicited by VGCC were as large and spread out as α9α10 transients (Figs. 3 and 6). This might indicate that, although smaller than in IHCs, Ca^2+^ transients through VGCC could be sufficient to evoke glutamate release onto type II afferent fibers. Moreover, these transients were further potentiated by RyR agonistic agents (Fig. 7). Whether RyRs mediating this effect are located in cisterns at synaptic (Grant et al., 2006; Lioudyno et al., 2004), or lateral wall locations (Grant et al., 2006; Ranum et al., 2019) is unknown. However, it suggests that CICR mechanisms can boost synaptic strength in contacts with type II afferents. CICR has been shown to modulate vesicle release and recruitment on afferent synapses of hair cells from frogs and turtles (Castellano-Muñoz et al., 2016; Lelli et al., 2003). Rod photoreceptors synapses are also modulated by this phenomenon, suggesting that CICR is not uncommon in ribbon synapses (Babai et al., 2010; Cadetti et al., 2006). In addition, ATP-induced IP3 mobilization and Ca^2+^ influx in the apical portion of isolated OHC, could further facilitate the Ca^2+^ spread down to synaptic sites (Ashmore & Ohmori, 1990; Mammano et al., 1999). This could explain the proposed function of type II neurons in pain sensation (Liu et al., 2015).

According to results in Figs. 4 and 7, SERCA pumps and RyR respond differently to afferent and efferent Ca^2+^ influx. One important question that arises is if, similarly to that described for immature IHCs (Moglie et al., 2018), OHCs cistern and buffering prevent Ca^2+^ spill-over from efferent to afferent synapses, particularly during sustained MOC activity. Both functional (Fig. 6E) and structural (Fuchs et al., 2014; Saito, 1990) evidence indicates that the diffusion interval between afferent and efferent synaptic locations is very short. This mechanism could operate in conjunction with VGCC to overcome an apparently weak synaptic drive in OHCs.

### Sorcin and Ca^2+^ regulation in OHCs

Transcriptome analysis has shown that gene products responsible for Ca^2+^ regulation, such as oncomod-ulin and sorcin, appear among the most highly represented mRNAs in OHCs (Li et al., 2018; Ranum et al., 2019; Shen et al., 2015). The high expression profile of oncomodulin agrees with previous reports showing a high concentration of the protein detected by immuno-EM (Hackney et al., 2005). Initially identified in drug-resistant cells, sorcin was later detected in cardiac myocytes, where it modulates CICR and excitation-contraction coupling in the heart (Colotti et al., 2014; Farrell et al., 2003; Lokuta et al., 1997; Meyers et al., 1995). Sorcin can also interact with, and regulate, other proteins responsible for Ca^2+^ homeostasis such as the Na+-Ca^2+^ exchanger (Zamparelli et al., 2010), L-type Ca^2+^ channels (Fowler et al., 2009; Meyers et al., 1998), and cisternal ATPases (Matsumoto et al., 2005).

The present work shows for the first time a function for sorcin in Ca^2+^ modulation at the synaptic pole of the OHCs. The addition of sorcin into the OHCs cytoplasm caused an increase in the resting cytoplasmic Ca^2+^ concentration (Fig. 5). One reasonable scenario could be that sorcin interacted with SERCA pumps located in sub-synaptic cisterns, inhibiting its action. However, this contrasts the reported role of sorcin on sarcoplasmic reticulum ATPases in the heart (Matsumoto et al., 2005). Another possibility would be that sorcin modulated Na+-Ca^2+^ exchangers as described for cardiomyocytes (Zamparelli et al., 2010). Interestingly, sorcin produced a reduction in the size of synaptic currents (calculated as integral, Q) during efferent stimulation (Fig. 5). In the same set of experiments Ca^2+^ transients were unaffected, implicating that sorcin does not affect α9α10 function, as these receptors are the sole reported Ca^2+^ source at this synapse. Since the recording conditions used for MOC stimulation experiments (Vh = −100 mV) were designed to maximize the Ca^2+^ driving force, part of the inward current triggered during synaptic events derives from SK2 channel activation. Thus, it is very likely that the sorcin mediated reduction of synaptic currents size is due to SK2 channels inhibition. One possible scenario is that sorcin operates as a sensor for Ca^2+^ influx through nicotinic receptors, curbing SK2 function to prevent over-inhibition by the MOC input.

The concentration of sorcin used in our experiments matches those in studies showing effects on RyR, Ca^2+^ channels, and Na+-Ca^2+^ exchangers (1-5 μM) (Farrell et al., 2003; Fowler et al., 2009; Lokuta et al., 1997; Zamparelli et al., 2010). Independently of its mechanism of action, our experiments indicate that sorcin is not merely a Ca^2+^ buffer. According to our experiments, the amount of additional buffer that sorcin represent in the electrode solution is negligible, < 0.5% (3 μM sorcin in a solution with 400 μM Fluo-4 and 500 μM EGTA), precluding a mere buffering effect. Moreover, the reported concentration of oncomodulin and other proteinaceous buffers in OHCs cytoplasm (in the mM range) (Hackney et al., 2005) would largely out compete the buffering capacity of a few μM concentration of sorcin. It was recently suggested that sorcin could operate as a ‘brake’ for Ca^2+^ spread originated in lateral walls cisterns of OHCs (Ranum et al., 2019), which implicates that sorcin would be a modulator of electromotility (Dallos et al., 1997; Frolenkov et al., 2000). Although further experiments are needed in order to decipher the role of sorcin in OHCs, the present results indicate a clear role for this protein in Ca^2+^ homeostasis.

## Methods

### Electrophysiological recordings from OHCs

Euthanasia and tissue extraction were carried out according to approved animal protocols of INGEBI Institutional Animal Care and Use Committee. Excised apical turns of 12- to 14-day-old mouse cochleas (Balb/c, either sex) were placed into a chamber on the stage of an upright microscope (Olympus BX51WI) and used within 2 hrs. OHCs were visualized on a monitor via a water immersion objective (60x), difference interference contrast optics and a CCD camera (Andor iXon 885). All recordings were performed at room temperature (22–25°C). Due to the short viability of the cochlear preparation and OHCs at this age, only one cell could be recorded per animal.

The cochlear preparation was superfused continuously at 2–3 ml/min with extracellular saline solution of an ionic composition similar to that of the perilymph (in mM): 144 NaCl, 5.8 KCl, 1.3 CaCl_2_, 0.7 NaH_2_PO_4_, 5.6 D-glucose, 10 HEPES buffer, 2 Pyruvate, 3 myo-inositol, pH 7.4. Working solutions containing different drugs were made up in this same saline and delivered through the perfusion system. Recording pipettes were fabricated from 1-mm borosilicate glass (WPI), with tip resistances of 6–8 MΩ. Series resistance errors were not compensated for.

For all experiments, the basic pipette solution was made from a 1.25X stock to reach a final concentration of (in mM): 95 KCl, 40 K-ascorbate, 5 HEPES, 2 pyruvate, 6 MgCl_2_, 5 Na_2_ATP, 10 Phosphocreatine-Na_2_, 0.5 EGTA and 0.4 Ca^2+^ indicator (Fluo-4), pH 7.2. To avoid variations in OHCs volume during experiments, pressure in the recording system was controlled with a digital manometer and kept within 5-9 cm H_2_0 range.

Heterologous expression and purification of human sorcin was performed as described in Meyers *et al*. (1995), with minimal adaptations. PET23d-Sorcin-wt was obtained from Dr. Gianni Colotti, was transformed into *Escherichia coli* BL21 (DE3) codon plus pLysS. Bacteria were growth in LB containing 5 mM CaCl2 until OD600 = 0.5, when 1 mM isopropylthiogalactoside (IPTG) was added and further incubated for 2 hours. Cells were harvested by centrifugation and washed with Lysis buffer (10 mM Tris-HCl, 10 mM NaCl, pH=7.5), followed by sonication (Sonication buffer: Lysis buffer with 1 mM DTT and antiproteases). Lysate was washed and resuspended in Sonication buffer plus 5mM MgCl_2_ and 0.2 μg DNAse (Fermentas). Following centrifugation, the supernatant was loaded into a Sep-Pak Accell Plus QMA cartridge (Waters), pre-equilibrated with Sonication buffer. Following a 5-step elution with 50, 150, 250, 400 and 500mM NaCl (2 ml each), SDS-PAGE revealed that most sorcin eluted in the 150 and 250 mM fractions. Samples were pooled in G2 dialysis cassettes (3500 MWCO, Thermo Scientific) and dialyzed twice for 12 hours at 4°C against 1 liter of (106.25 KCl, 6.25 HEPES, 2.5 Pyruvate and 7.5 MgCl_2_). After dialysis, samples were concentrated using Vivaspin devices (3000 MWCO, Cytiva) and protein concentration was estimated by the Bradford assay.

Stock solutions of dantrolene, thapsigargin and ryanodine (both at 1 and 100 μM) were prepared in DMSO and added to the intracellular solution, such that the concentration of DMSO in the pipette solution was 0.5 % v/v in every case. All salts and drugs were acquired from SIGMA except for ryanodine, dantrolene and thapsigargin which were purchased from Tocris.

Efferent synaptic currents were evoked by unipolar electrical stimulation of the MOC efferent axons as described previously (Ballestero et al., 2011; Goutman et al., 2005). Briefly, the electrical stimulus was delivered via a 20 to 80 μm-diameter glass pipette which position was adjusted until postsynaptic currents in OHCs were consistently activated. An electrically isolated constant current source (model DS3, Digitimer) was triggered via the data-acquisition computer to generate pulses of 40 to 220 μA, 1 msec width. Further increasing stimulation intensity, or duration, did not produce any rise in release probability (or reduction in failure rate). Solutions containing ACh were applied by a gravity-fed multichannel glass pipette (150 μm tip diameter).

Electrophysiological recordings were performed using a Multiclamp 700B amplifier (Molecular Devices), low-pass filtered at 6 kHz and digitized at 50 kHz via a National Instruments board. Data was acquired using WinWCP (J. Dempster, University of Strathclyde). To maximize Ca^2+^ driving force during imaging experiments, OHCs were voltage clamped at −100 mV, but only during a brief period of time when stimulation was applied and synaptic responses were recorded (650 msec in paired pulse experiments, and 2 sec in trains). Otherwise, cells were held at −40 mV. Electric shocks to MOC fibers were separated by intervals of 5 sec in paired pulse experiments, and 30 sec for trains. Recordings were analyzed with custom-written routines in IgorPro 6.37 (Wavemetrics). IPSCs were evaluated as successfully activated when met a double criteria: i) the amplitude of the current right after the stimulation artifact was > 3x S.D. of the basal current, and ii) the integral of the IPSC was > 0.3 pC, which is the average integral of a single synaptic event (Ballestero et al., 2011).

### Ca^2+^ Imaging Experiments

Ca^2+^ indicators were included in the patch-pipettes (at the concentration indicated before) allowing the diffusion into the cells. The preparation was illuminated with a blue LED system (Tolket, Argentina) and images were acquired using an Andor iXon 885 camera controlled through a Till Photonics interface system. The focal plane was set close to the basal pole of OHCs where synapses are found. The signal-to-noise ratio was improved with an on-chip binning of 4×4, giving a resolution of 0.533 μm per pixel with the 60X water immersion objective. The image size was set to 50×50 pixels which allowed an acquisition rate of 140 frames/sec. Image acquisition started 5 min after whole-cell break in to ensure the proper dialysis of the cell content and lasted up to 45 min. Images were analyzed with custom-written routines in IgorPro 6.37 (Wavemetrics).

A time lapse consisting of 250 images were taken for each experiment. An averaged image in each time lapse was used to determine the edge of the cell by an automatic thresholding algorithm. Within the cell borders, a donut-shaped mask covering the cell’s cytoplasm was defined comprising 40 to 90% of the maximal fluorescence signal. The mask was divided in 24 radial regions of interest (ROIs) with its center set at the maximal intensity pixel of the cell. Fluorescence intensity was measured in every ROI for each time frame. Two criteria were used to determine that a successful synaptic Ca^2+^ event occurred at a particular ROI: i) a fluorescence peak was identified right after the MOC stimulus, with 2.5x higher amplitude than the standard deviation of the baseline fluorescence; and ii) the area under the curve (fluorescence trace) was larger than 0.11 (A. U. * sec). Finally, those ROIs that exhibited a consistent pattern of activation were selected as hotspots and used for further analysis of the fluorescence signal. Photobleaching was corrected for long acquisition protocols by fitting a line between pre-stimulus baseline and final fluorescence.

To determine the spread of the fluorescence change across the OHC cytoplasm, the response image when the fluorescence signal peaked was normalized to pre-stimulus fluorescence. Then, it was thresh-olded by fitting a bimodal distribution to the image histogram and the area and center of mass of the resulting mask calculated. Signal spread area was divided by the cell total area for comparison. The center of mass, which takes into account not only the area of the signal but also the fluorescence intensity in each point, was used to determine the location of the Ca^2+^ entry sites (both MOC and VGCC), and thus the distance between them.

### Statistical Analysis

Data are presented as group means ± standard error (SEM) and were analyzed with Infostat (Universidad Nacional de Córdoba). The Mann-Whitney test was used to perform comparisons between two groups and Kruskal-Wallis one-way analysis of variance followed by Conover’s test for comparisons between multiple groups. Wilcoxon signed-rank and Friedman’s test were used for comparisons between two and multiple paired samples, respectively. Non-parametric statistics were preferred given our sample sizes and the impossibility of testing the assumptions of parametric procedures. F-test was used to compare models fitting in Figure 5C. Differences between samples were considered significant when p<0.05. Effect sizes were calculated for Figs. 3, 4, 5 and 7 using R Statistical Software (RRID:SCR_001905) and the statsExpressions package. Values obtained in each case and the parameter of the effect size implemented are informed in a supplementary document.

## Supporting information

Supplemental information on Effect Sizes of statistical tests

Point by Point response to reviewers

## Acknowledgements

Maryline Beurg for advice on the use of ascorbic acid in the intracellular solution, and Joe Santos-Sacchi on OHCs recordings. Gianni Colotti for kindly sharing the plasmid vector for sorcin production, and J. Dempster for the use of WinWCP. Paul A. Fuchs for comments on the manuscript.

