## Supplemental information on Effect Sizes of statistical tests for "Synaptic contributions to cochlear outer hair cell Ca^2+^ homeostasis"

**Figure 3**

**Test**: Friedman's rank sum test
**Effect size**: Kendall's *W*

| Panel | Variable | $W_{Kendall}$ | $k$* | $W_{Kendal}$** | $k$ |
|---|---|---|---|---|---|
| C | $\Delta F/F_0$ | 0.706 | 3 | 0.263 | 4 |
| D | Q | 0.937 | 3 | | |
| F | Area Signal | 0.762 | 3 | 0.692 | 4 |
| H | $\Delta F/F_0$ (Variable Stim Time) | 0.5 | 6 | | |

*$k$ = number of treatments
**Effect size calculated taking into account the depolarization protocols

**Figure 4**

**Test**: Wilcoxon signed-rank test
**Effect size**: $r = Z/\sqrt{Npairs}$

| Panel | Variable | Protocol | $r$ |
|---|---|---|---|
| B | $\Delta F$ (Baseline) | | 0.898 |
| | $\Delta F$ | 20 Hz | 0.301 |
| | $\Delta F$ | 40 Hz | 0.641 |
| | $\Delta F$ | 80 Hz | 0.903 |
| C | Q | 20 Hz | 0.541 |
| | Q | 40 Hz | 0.3 |
| | Q | 80 Hz | 0.541 |
| D | FWHM | 20 Hz | 0.0604 |
| | FWHM | 40 Hz | 0.812 |
| | FWHM | 80 Hz | 0.903 |

**Test**: Kruskal–Wallis test
**Effect size**: $\epsilon^2$

| Panel | Variable | Protocol | $\epsilon^2$ |
|---|---|---|---|
| F | $\Delta F$ (Baseline) | | 0.127 |
| G | FWHM | 20 Hz | 0.0418 |
| | FWHM | 40 Hz | 0.04 |
| | FWHM | 80 Hz | 0.087 |
| H | $\Delta F/F_0$ | 20 Hz | 0.0341 |
| | $\Delta F/F_0$ | 40 Hz | 0.158 |
| | $\Delta F/F_0$ | 80 Hz | 0.125 |
| | Q | 20 Hz | 0.0868 |
| | Q | 40 Hz | 0.0375 |
| | Q | 80 Hz | 0.139 |

**Figure 5**

**Test**: Two-sample Mann–Whitney *U* Test
**Effect size**: $r = Z/\sqrt{Npairs}$

| Panel | Variable | Protocol | *r* |
|---|---|---|---|
| B | ΔF (Baseline) | | 0.62 |
| C | ΔF | 20 Hz | 0.484 |
| | ΔF | 40 Hz | 0.0345 |
| | ΔF | 80 Hz | 0.069 |
| D | Q | 20 Hz | 0.449 |
| | Q | 40 Hz | 0.345 |
| | Q | 80 Hz | 0.585 |

**Figure 7**

**Test**: Wilcoxon signed-rank test
**Effect size**: $r = Z/\sqrt{Npairs}$

| Panel | Variable | Protocol | $r$ |
|---|---|---|---|
| A | $\Delta F$ | | 0.301 |

**Test**: Kruskal–Wallis test
**Effect size**: $\epsilon^2$

| Panel | Variable | Protocol | $\epsilon^2$ |
|---|---|---|---|
| B | $\Delta F/F_0$ | | 0.246 |

**Test**: Two-sample Mann–Whitney *U* Test
**Effect size**: $r = Z/\sqrt{Npairs}$

| Panel | Variable | Protocol | $r$ |
|---|---|---|---|
| C | $\Delta F$ | | 0.345 |
