## Supplementary material for "Synaptic contributions to cochlear outer hair cell Ca^2+^ homeostasis": Point by Point response to reviewers

**Reviewer #1 (General assessment and major comments (Required)):**

**This manuscript describes convincing measurements of cytoplasmic $Ca^{2+}$ signals attributable to voltage gated $Ca^{2+}$ channels and efferent nAChR channels. These channel coexist on the basolateral surface of OHCs and may, with the MET channels, contribute to OHC $Ca^{2+}$ homeostasis. The main conclusions are that the two channel types are differentially modulated, ryanodine receptor action potentiating VGCC but not efferents, which disagrees with previous claims (Lioudyno et al 2004); and efferent responses were reduced by sorcin, a $Ca^{2+}$-binding protein recently localized to OHCs, and known inhibitor of ryanodine receptors. Neither the concentration nor exact mechanism of sorcin's action was determined.**
**Specific points:**
**1. In the fluorescent images in the various figures, the image orientation is unclear- is it a radial or a transverse view? It would help if in some figures, a representation of an OHC, either as Nomarski image or as a drawing, can accompany the fluorescent image.**

This is an excellent suggestion by the reviewer. A drawing is now included in Fig. 1A

**2. L126. in describing $Ca^{2+}$ spread, especially with long stimulation, there is concern that the high affinity dye Fluo4 will saturate. This should be discussed. I would not have used this dye - preferred a lower-affinity dye such as Fluo5.**

We regret that this issue was not sufficiently clarified in our original version of the manuscript. It should firstly indicated that a high affinity indicator was selected because of the small $Ca^{2+}$ influx generated during physiological $\alpha9\alpha10$ activation, that was on the verge of our detection threshold. A lower affinity dye would not be able to resolve those $Ca^{2+}$ transients. This important observation is now clarified on line 94 on Results:

*"$\alpha9\alpha10$ receptors are highly $Ca^{2+}$ permeable, but due to the much higher concentration of $Na^+$ and $K^+$ in the extracellular solution (150 mM together vs 1.3 mM $Ca^{2+}$, see Methods), the majority of the current allowed in physiological conditions is due to monovalent cations (Weisstaub et al., 2002). To circumvent the small $Ca^{2+}$ influx, we adopted two technical measures. Firstly, a high affinity $Ca^{2+}$ indicator, such as Fluo-4, was used to detect synaptic $Ca^{2+}$ transients. And secondly, to maximize the $Ca^{2+}$ driving force, OHCs were transiently voltage-clamped at -100 mV..."*

Reviewer #1 also brought up the issue of saturation of the $Ca^{2+}$ dye. We believe that it is very unlikely that Fluo-4 is saturated in experiments included in Figs. 3-7 considering that $Ca^{2+}$ signals detected with the same indicator (and also repeating all meaningful conditions) but using ACh local application (Fig. 1) were ~20 times larger (compared to 80 Hz trains). The integral of the ACh response was >30 times larger than 80 Hz trains. It is likely that a high affinity indicator such as Fluo-4 is saturated in experiments with large, and sustained, $Ca^{2+}$ influx as those obtained in ACh applications. But the currents are significantly smaller, and briefer, during trains of MOC stimulations, indicating that Fluo-4 is within its dynamic range of $Ca^{2+}$ sensitivity. Moreover, the linear correlation between $\Delta F/F$ and charge in Fig. 3E points in the same direction.
To clarify this problem, a sentence was included in Results, line 143:

*"Comparing $Ca^{2+}$ signals in Figs. 1 and 3 it becomes clear that synaptic activation of $\alpha9\alpha10$ receptors during trains is (at least) ~20 times smaller than that obtained with locally applied ACh. This could be due to the rather low release probability of MOC terminals (Ballestero et al., 2011; Vattino et al., 2020), that our stimulation paradigm typically activates a single axon (see Methods), and that activation by ACh does not rely of vesicle availability. This drastic difference in measurable $\Delta F/F_0$ amplitude between ACh application and electrical stimulation also suggests that the $Ca^{2+}$ indicator used in these experiments, Fluo-4, were not saturated during synaptic trains. The tight correlation between $\Delta F/F$ and charge in Fig. 3E further supports this conclusion."*

And also in the Discussion section, line 271:
*"Synaptic $Ca^{2+}$ signals of both origins were largely smaller than those obtained by local perfusion of a high ACh concentration (compare Figs. 1, 3 and 6), suggesting that the $Ca^{2+}$ indicator used in these experiments (Fluo-4) was not saturated during physiological activation. The selection of the indicator was based upon the signal to noise ratio obtained in experiments that sought to detect $Ca^{2+}$ influx through the highly $Ca^{2+}$ permeable $\alpha9\alpha10$ receptors, which actually allows predominantly $Na^+$ through when activated in physiological conditions (140 mM $Na^+$ and 1.3 mM $Ca^{2+}$, as in our experiments) (Moglie et al., 2018; Weisstaub et al., 2002)."*

**3. Fig. 3F and L:124. Express the spread in absolute units rather than percent of OHC diameter. I assume the conclusion (not stated) is that the $Ca^{2+}$ rise is not confined to the sub-cisternal space but spreads throughout the cell. Why does it not activate release of the afferent neurotransmitter? A point not mentioned is that the efferent SK2 and BK channels are distributed along the lateral membrane.**

Expressing the $Ca^{2+}$ signal spread in absolute units is a good suggestion. But we favor the % of the imaged cytoplasmic space because it provides an idea of how spread out is $Ca^{2+}$ influx in the context of the entire cell, and thus, it conveys biological meaning.
The reviewer is right in that the imaged $Ca^{2+}$ may escape the sub-cisternal space. This fact was mentioned in Discussion, line 302 (transcribed below) together with the possibility that this $Ca^{2+}$ rise could activate release on the afferent synapse (line 343).

line 302:

*"Evidence from Figure 4, using train stimulation to MOC fibers, suggests that cisterns are responsible not only for speeding up the decay of efferent $Ca^{2+}$ but also limiting its spread. However, taking into account the volume of this small domain and the spread of the $Ca^{2+}$ transients (Fig. 3F), it is very likely that $Ca^{2+}$ signals evoked at high frequency MOC stimulation result, at least partially, from $Ca^{2+}$ that escaped into the cytoplasm. A circumstance that may differ in conditions with more intact intracellular buffers concentrations."*

line 343:

*"One important question that arises is if, similarly to that described for immature IHCs (Moglie et al., 2018), OHCs cistern and buffering prevent $Ca^{2+}$ spill-over from efferent to afferent synapses, particularly during sustained MOC activity. Both functional (Fig. 6E) and structural (Fuchs et al., 2014; Saito, 1990) evidence indicates that the diffusion interval between*

*afferent and efferent synaptic locations is very short. This mechanism could operate in conjunction with VGCC to overcome an apparently weak synaptic drive in OHCs."*

To the best of our knowledge, SK and BK channels are located at synaptic sites (Wersinger et al., 2010 Plos One).

**4. Fig. 6F. Ideally the spread of $Ca^{2+}$ signals at the peak should be presented as (overlapping) Gaussians for the two sources. The significance of the 3.7 um separation (L319) between the sources needs some context.**

The Gaussian representation of the $Ca^{2+}$ spread would be a possible way to analyze the data. But we think that it imposes a strong assumption on the shape of the spread that may not be met by our signals. Particularly considering the limits imposed by the cell shape. We took an alternative approach that did not requires such assumption. We calculated the center of mass of the hotspot area, and used it as a coordinate to measure distances between afferent and efferent $Ca^{2+}$. This description is included in Methods (now improved, see below), and we now introduced a reference to this section in results.
Line 469 Methods:

*"The center of mass, which takes into account not only the area of the signal but also the fluorescence intensity in each point, was used to determine the location of the $Ca^{2+}$ entry sites (both MOC and VGCC), and thus the distance between them."*

We also discussed the physiological consequences of this tight localization (see line 345):

*"... Both functional (Fig. 6E) and structural (Fuchs et al. 2014; Saito, 1990) evidence indicates that the diffusion interval between afferent and efferent synaptic locations is very short. This mechanism could operate in conjunction with VGCC to overcome an apparently weak synaptic drive in OHCs."*

**5. L169. State explicitly that the ryanodine results disagree with (Lioudyno et al 2004).**

A sentence making this fact more explicit is now included in Discussion (line 312):

"*This result contrasts with previous observations in OHCs in which the role of RyR was evaluated, not directly on Ca2+ transients as we did here, but by measuring efferent synaptic activation (Lioudyno et al., 2004)."*

**6. L177. Refer to Corey's Shield database (Scheffer et al 2015) who first reported the presence of sorcin mRNA in OHCs.**

Corrected.

**7. L178. A concern is over the physiological significance of the sorcin effects. Sorcin is a $Ca^{2+}$ binding protein that if present at high concentrations could supplement oncomodulin in addition to inhibiting RyRs. Can the authors determine the sorcin concentration in OHC cytoplasm? In addition it seems strange that the reported effect to sorcin is to inhibit RyRs so limiting the temporal spread of the CICR, but the present results suggest Can the authors clarify these problems.**

In our set of experiments we intended to provide a characterization of sorcin effects on $Ca^{2+}$ homeostasis of OHC, using the concentration of the protein that has shown a strong effect on RyRs and in cardiac myocytes. Looking at isolated RyRs activity, Lokuta and colleagues found that the Po decreased with the exposure to sorcin, with an IC50 of 0.5 μM (Lokuta et al., 1997). Meanwhile, in intact cardiac myocytes studies, sorcin at 1 to 5 μM showed a strong inhibition of voltage activated $Ca^{2+}$ transients and other $Ca^{2+}$ mediated processes. Similarly, it was also shown that sorcin can effectively modulate L-type $Ca^{2+}$ channels (Fowler et al., 2009), cisternal ATPases (Matsumoto et al., 2005) and $Na^+$-$Ca^{2+}$ exchangers (Zamparelli et al., 2010) within the same range of sorcin concentrations. In our experiments we only added 3 μM sorcin to an electrode solution with 400 μM fluo-4 and 500 μM EGTA. Thus, sorcin represented only 0.3% of additional $Ca^{2+}$ buffer. In addition, our results suggests that sorcin does not operate solely as a $Ca^{2+}$ buffer: i) adding extra buffer would only produce a reduction in resting $Ca^{2+}$, not the opposite (Fig. 5B); ii) $Ca^{2+}$ transients elicited by MOC synapse activation would be reduced (as in Moglie et al., 2018 Fig. S7), instead of no change (Fig. 5D).

On the other hand, it has been shown that millimolar concentrations of oncomodulin are expressed in OHCs cytoplasm (Hackney et al., 2005). Therefore, if one assumes that sorcin effects in OHCs are produced by a few μM concentration of this protein, the buffering capacity of OHCs would not be affected.

Some of these arguments were included in the new version of Discussion, line 375:

*"The concentration of sorcin used in our experiments matches those in studies showing effects on RyR, $Ca^{2+}$ channels, and $Na^+$-$Ca^{2+}$ exchangers (1-5 μM) (Farrell et al., 2003; Fowler et al., 2009; Lokuta et al., 1997; Zamparelli et al., 2010). Independently of the mechanism of action, which should be investigated further, our experiments indicate that sorcin is not merely a $Ca^{2+}$ buffer. According to our experiments, the amount of additional buffer that sorcin represent in the electrode solution is negligible, <1% (3 μM sorcin in a solution with 400 μM Fluo-4 and 500 μM EGTA). None of the effects observed in Figs. 5 and 7 could be explained by that. Moreover, the reported concentration of oncomodulin and other proteinaceous buffers in OHCs cytoplasm (in the mM range) (Hackney et al., 2005) would largely out compete the buffering capacity of a few μM concentration of sorcin."*

**8. L199-204. The authors could have resolved whether sorcin affected SK2 channels by (briefly) switching to -40 holding potential where the nAChR and SK2 currents would be of opposite polarity.**

We agree that this would be an alternative way of looking at SK2 inhibition. Our purpose was to evaluate both $Ca^{2+}$ transients and synaptic currents simultaneously to provide a comprehensive view of the effect of sorcin on synaptic function.

**9. L350 omit 'novel' Sorcin is not a novel protein having been described in the 1990's**
Done

**Reviewer #2 (General assessment and major comments (Required)):**

**In this study, the group of Juan Goutman investigated $Ca^{2+}$ signaling in immature cochlear outer hair cells (OHCs). The work focuses on the basolateral compartment analyzing $Ca^{2+}$ signals mediated by afferent ribbon-type active zones and by efferent synapses. $Ca^{2+}$ influx at the ribbon-type active zones is substantial, which is in keeping with the large ribbons found in OHCs. The authors show that it can be potentiated by ryanodine which indicates an interesting interplay between voltage-gated $Ca^{2+}$ influx and ryanodine receptor mediated $Ca^{2+}$ release from internal stores. Finally, adding recombinant sorcin, a $Ca^{2+}$ binding protein prominently expressed in cardiomyocytes to the patch-pipette modulated the basal $[Ca^{2+}]i$ and efferent $Ca^{2+}$ signalling in OHCs. The authors provide characterization of efferent and afferent $Ca^{2+}$ signals. An interesting paper and suitable for publication at eLife but a number issues that keep me from recommending publication in its present form.**

**Novelty:**

**The approach taken and some of the conclusions is similar to what the group presented for immature inner hair cells, that also feature afferent and efferent synapses in close proximity and with functional interaction. This absolutely reasonable to do but presents an extension of the same concept to a related cell type.**

We thank the reviewer for the comments

**Relevance for understanding OHC function in the mature cochlea:**

**The authors have performed experiments on organs of Corti from mice at postnatal days 12-14. This is around the onset of hearing in mice and represents a time window during which substantial changes have been shown to occur. Figures 4 and 5 of Hackney et al., JN2005 show that the cytosolic abundance $Ca^{2+}$ binding proteins parvalbumin a, parvalbumin ß, and calretinin changes dramatically around this stage of development. Hence, the presented data should not be taken to conclude on the situation in the mature cochlea.**

We tried to keep our study within close limits of interpretation, by putting results into context, and pointing out at technical limitations, including buffering concentration. In this respect, in line 304, we included the following sentence:

*"However, taking into account the volume of this small domain and the spread of the $Ca^{2+}$ transients (Fig. 3F), it is very likely that $Ca^{2+}$ signals evoked at high frequency MOC stimulation result, at least partially, from $Ca^{2+}$ that escaped into the cytoplasm. A circumstance that may differ in conditions with more intact intracellular buffers concentrations."*

And now, we included the following sentence to address the developmental issue, on line 307:

*“Due to the fragility of the tissue, as well as the instability of OHCs recordings from older animals, we decided to circumscribe our experiments within the P12-P14 age range. Therefore, some of our conclusions may differ in more mature OHCs.”*

**Statistical data basis/sample size:**

**Analyzing highly variable Ca2+ signals in hair cells poses the challenge of capturing the underlying distribution by sufficient sample size. Several experiments in the present study fall short in acquiring such sample size.**

As the reviewer points out, these are challenging experiments and we tried to make as many repetitions as possible, particularly in those experiments that could bring the more intense debate (sorcin and ryanodine). The analysis of effect sizes for each statistical test is also included for the reviewers’ evaluation as supplemental information.

**Role of sorcin:**

**I highly recommend the authors to provide own sorcin immunohistochemistry. Perfusion of the cytosol with recombinant Ca2+ binding proteins is expected to affect Ca2+ signalling (reducing amplitude and spread) and in a way similar to the addition of synthetic Ca2+ chelators. With 3 µM of recombinant protein, it seems difficult to achieve a sizable effect (even when considering fully functional multiple EF-hands. In the present study, a non-significant trend towards a reduced amplitude of afferent Ca2+ signals was observed during whole-cell patch clamp with sorcin (molar concentration should be provided). The relevance of sorcin function for OHC function remains to be studied by deleting sorcin expression in OHCs and performing comparative perforated-patch recordings from sorcin-deficient mice or siRNA knock-down.**

We are thankful to the reviewer for the suggestion of very interesting experiments. Looking at the functional phenotype of sorcin KO mice would be a fundamental step towards understanding the role of this $Ca^{2+}$ binding protein in the auditory system. However, it is beyond the scope of the current study.
Ranum and colleagues (2019) have provided inmunohistochemistry anti sorcin in OHCs, confirming the presence of the protein as well. We are currently planning a more comprehensive study in which the sub-cellular location of sorcin will be evaluated together with markers for cisterns, and also investigating changes in the protein partitioning depending on $Ca^{2+}$ concentrations. These experiments are beyond the scope of our current study.

**Specific comments:**

**Mention species in title and/or abstract**

Done. The abstract now reads (line 17):

*“Using functional imaging in rodent OHCs, we report that these two $Ca^{2+}$ entry sites are closely positioned, but present different regulation mechanisms.”*

**What is meant by "we found that VGCC Ca2+ signals are larger than expected" please disambiguate or remove?**

We agree with the reviewer that this sentence is not clear. As suggested, it was modified (line 19):

*"Considered a weak drive for transmitter release, we unexpectedly found that VGCC $Ca^{2+}$ signals are comparable in size to those by $\alpha 9 \alpha 10$ and can be potentiated by ryanodine receptors."*

**Also consider replacing "VGCC Ca2+ signals" by afferent or presynaptic Ca2+ signals, as the proposed CICR contribution indicates a more complex origin of Ca2+ contributing to these signals.**

With 'VGCC $Ca^{2+}$ signals' we refer to those generated by activation of VGCC, irrespective of whether they are further modulated by RyR. We believe that this is a more general way to refer to these signals within the context of an abstract.

**Line 56: "we found that Ca2+ signals from VGCC are unexpectedly large," see my comment above**

In contrast to the commented sentence in Abstract, this one in Introduction was preceded by contextual sentences indicating that VGCC in OHCs were very small and considered unfit for triggering synaptic release. Our $Ca^{2+}$ imaging results show that the corresponding $Ca^{2+}$ signals are actually 'unexpectedly large'.

**Line 57 and throughout: consider clarifying that you refer to signal amplitude not spatial extent of the signal (perhaps replace size by dF/F0 or amplitude)**

Suggestion taken. Line 57 now reads:
"*...comparable in amplitude with $\alpha 9 \alpha 10$ transients*"

**Line 61: "control Ca2+-based excitation-contraction coupling in cardiomyocytes"?**

This sentence was corrected with a more explicit version (line 60):
"*It has been shown that sorcin controls excitation-contraction coupling in cardiomyocytes, by modulating the activity of RyRs, L-type VGCC and other proteins involved in $Ca^{2+}$ homeostasis (Colotti et al., 2014)*"

**Line 62: "among the most differentially expressed genes in OHCs" this statement is not useful without mentioning the cells used for the comparison**

Sentence corrected. Now it reads (line 63):
"More recently, it has been reported that sorcin is among the most differentially expressed genes in OHCs (compared to IHC and supporting cells) (Li et al., 2019; Ranum et al., 2019; Shen et al., 2015), although its role remained hypothetical."

**Line 64: "Thus, the present results shed light into Ca2+ homeostasis in the hair cells**

**involved in sound amplification at the cochlea, and unveil a role for the novel protein sorcin."**
**I don't think so, please see major concerns.**
In this study we characterized how synaptic $Ca^{2+}$ dynamics were affected by the action of multiple players in the network of $Ca^{2+}$ control in OHCs. There was previous evidence indicating that RyR, SERCA pumps and even the $Na^+$-$Ca^{2+}$ exchangers could modulate intracellular $Ca^{2+}$ in OHCs, but we now provide a direct assessment of how these molecules affect both efferent and afferent synaptic $Ca^{2+}$. In addition, we also provide experimental evidence indicating that sorcin should be added to this list of modulators of intracellular $Ca^{2+}$ concentration in OHCs. We believe that a role for sorcin is clearly shown in Figs. 5 and 7. Our plan is to further investigate mechanisms of action in the near future.

**Line 70 and following: I think this first section is mainly confirmatory (work by the Mammano lab and others) and hence might better serve as supplementary information. Please add whether the data points in C-E correspond to cells and singe trials or represent average responses of each OHC.**

We presume that this sentence refers to Figure 1 in our manuscript. We believe that this figure helps illustrate the imaging protocol developed in our study, and more importantly, on the upper bound of the $Ca^{2+}$ signal activated by $\alpha9\alpha10$ receptors. As indicated in the response to other reviewers, we believe that a misinterpretation occurred regarding the dynamic range of the $Ca^{2+}$ indicator that was selected for this study. Figure 1 is instrumental in this context (see responses to reviewers #1 and #3). Nonetheless, we agree that previous work by Mammano and others are critical, and thus, were cited accordingly (e.g. Evans et al 2000; Frolenkov et al., 2000; Lelli et al., 2003; Mammano et al., 1999).

**Line 88: So, do you assume that the hotspot corresponds to a single efferent OHC synapse being activated?**

Across experiments, a single hotspot could be detected in each OHCs. Considering the high activation threshold of MOC synapses in our hands (compared to IHC in Moglie et al., 2018), we reasoned that the most likely scenario would be that a single synapse was activated in each OHCs.

**Line 97:Was this averaging including the failures? If not the example shown in Fig. 2B does not really seem representative? Consider adding a note relating the dF/F0 for ACh and efferent transmission: 2 orders of magnitude difference. Also please reflect on finding failing $Ca^{2+}$ signals despite successful IPSC.**

All these observations are important and require clarification in the text. Regarding average values on line 97, they actually exclude failures, and this fact is now indicated in line 107:

*"In paired pulse protocols, an average $Ca^{2+}$ signal of 2.7 ± 0.3 %$\Delta F/F_0$ (n = 6 cells) was obtained, whereas the integral of the synaptic currents was 7.4 ± 0.8 pC, excluding synaptic failures for the calculation of both averages (n = 6) (Fig. 2C and D)"*

The following line was also added to line 113, to address the finding of a small group of successful IPSCs with no detectable $Ca^{2+}$ signals:

*“Note that failures in imaging events coincided with the smallest IPSCs (black symbols in Fig. 2E).”*

We favor the inclusion in Fig. 2B of $Ca^{2+}$ transient traces corresponding to synaptic failures, because it provides an idea of how different successful events differ from failures.

In relation to the following comment: ‘**Consider adding a note relating the ΔF/F0 for ACh and efferent transmission: 2 orders of magnitude differences**’, we actually added new paragraphs in Results and Discussion referring to this fact. As indicated in the response to other reviewers, this striking difference argues against a saturation of our $Ca^{2+}$ indicator in experiments with trains of stimuli.
On line 143 of Results, it now reads:

*“Comparing $Ca^{2+}$ signals in Figs. 1 and 3 it becomes clear that synaptic activation of $\alpha 9 \alpha 10$ receptors during trains is (at least) ~20 times smaller than that obtained with locally applied ACh. This could be due to the rather low release probability of MOC terminals (Ballestero et al., 2011; Vattino et al., 2020), that our stimulation paradigm typically activates a single axon (see Methods), and that activation by ACh does not rely of vesicle availability. This drastic difference in measurable $\Delta F/F_0$ amplitude between ACh application and electrical stimulation also suggests that the $Ca^{2+}$ indicator used in these experiments, Fluo-4, were not saturated during synaptic trains. The tight correlation between $\Delta F/F$ and charge in Fig. 3E further supports this conclusion.”*

And on line 271 in Discussion:

*“Synaptic $Ca^{2+}$ signals of both origins were largely smaller than those obtained by local perfusion of a high ACh concentration (compare Figs. 1, 3 and 6), suggesting that the $Ca^{2+}$ indicator used in these experiments (Fluo-4) was not saturated during physiological activation. The selection of the indicator was based upon the signal to noise ratio obtained in experiments that sought to detect $Ca^{2+}$ influx through the highly $Ca^{2+}$ permeable $\alpha 9 \alpha 10$ receptors, which actually allows predominantly $Na^+$ through when activated in physiological conditions (140 mM $Na^+$ and 1.3 mM $Ca^{2+}$, as in our experiments) (Moglie et al., 2018; Weisstaub et al., 2002)”*

**Legend to fig. 2 should mention the imaging approach used here. Please add whether the data points in C-E correspond to cells and singe trials or represent average responses of each OHC. "during double-pulse"**

We believe that reviewer #2 is right in that a consistent legend should include the imaging approach, just like it was done in Fig. 1. Instead, we decided to remove details from Fig. 1 legend, as all experiments were performed, and analyzed, with similar procedures and all the information is included in Methods.

**Line 102: Consider to move this explanation up to where you introduce the experiment.**

We thank the reviewer for this suggestion, which is actually very useful. Line 91 in Results now reads:

*"Representative traces of fluorescence changes at the brightest ROI are shown in the bottom panel of Figure 2B (red traces), whereas the corresponding synaptic currents recorded in the same trials are included in the top panel (black traces). In order to maximize the $Ca^{2+}$ driving force, OHCs were transiently voltage-clamped at -100 mV in these experiments (for the duration of the synaptic response, otherwise, at -40 mV. See Methods) such that inward currents were due to the activation of both $\alpha 9\alpha 10$ receptors and SK2 channels ($K^+$ equilibrium potential: ~-82 mV)."*

**Line 117: A methods section detailing the statistical analysis is missing completely. How was the use of a non-parametric test (Friedman's test) justified: i.e. how was normality tested?**

We regret having missed the importance of this section. Non-parametric statistics was used due to the impossibility of testing the assumptions of parametric statistics. Our mistake was corrected in the new version of the manuscript, as it read now in Methods, line 473:

"*Statistical Analysis*
*Data are presented as group means ± standard error (SEM) and were analyzed with Infostat (Universidad Nacional de Córdoba). The Mann-Whitney test was used to perform comparisons between two groups and Kruskal-Wallis one-way analysis of variance followed by Conover's test for comparisons between multiple groups. Wilcoxon signed-rank and Friedman's test were used for comparisons between two and multiple paired samples, respectively. Non-parametric statistics were preferred given our sample sizes and the impossibility of testing the assumptions of parametric procedures. F-test was used to compare models fitting in Figure 5C. Differences between samples were considered significant when p<0.05. Effect sizes were calculated for Figs. 3, 4, 5 and 7 using R Statistical Software (RRID:SCR_001905) and the statsExpressions package. Values obtained in each case and the parameter of the effect size implemented are informed in a supplementary document.*"

**Line 122: "localized Ca2+ rise with a measurable spread which accounted for 31 {plus minus} 5 % of the area corresponding to the imaged OHC area." How was "measurable spread" defined?**

A detailed explanation for how spread was estimated is included in Methods, line 466:

*"To determine the spread of the fluorescence change across the OHC cytoplasm, the response image when the fluorescence signal peaked was normalized to pre-stimulus fluorescence. Then, it was thresholded by fitting a bimodal distribution to the image histogram and the area and center of mass of the resulting mask calculated. Signal spread area was divided by the cell total area for comparison. "*

**Line 128: The maximal Ca2+ signal with 80 Hz stimulation of efferent synapses is still an order of magnitude lower than that found with ACh. The authors suggest that the Ca2+ rise is limited by SERCA pumps, but do they assume, indeed, that this clearance mechanism is not at work during ACh application?**

This is an interesting point. We believe that during synaptic activation of $\alpha 9\alpha 10$, SERCA and other mechanisms of $Ca^{2+}$ control, are in place; even during long stimulation trains, at rather high frequencies, such as 80 Hz. But with ACh applications, $Ca^{2+}$ influx is massive, producing a huge concentration rise during seconds in OHCs cytoplasm. It is possible that SERCA pumps are unable to handle this $Ca^{2+}$ rise, and that little differences would be observed in the absence of SERCA extrusion. As indicated, there is substantial difference between synaptic activation of $\alpha9\alpha10$, and that generated by ACh applications.

**Line 141: How sure can the authors be that this cytosolic Ca2+ rise does not result from a store-depletion related Ca2+ entry?**

Regardless of the origin of this $Ca^{2+}$ rise, our point is that when SERCA pumps are inhibited cytoplasmic $Ca^{2+}$ increases. This reflects that there is a basal extrusion rate operating in OHCs.

**Line 155: I recommend keeping the order from 20-80 Hz as above and below to make reading easier.**

Suggestion taken.

**Line 178: How confident can we be that the recombinant sorcin was Ca2+ free, in other words, could the elevated basal Ca2+ simply reflect preloading of sorcin?**

As indicated in Methods, sorcin was purified from bacteria using Sep-Pak Accell Plus QMA cartridges and washed 5 times. Eluted fractions containing sorcin protein were dialyzed twice for 12 hours against 1 liter of a solution that was prepared with the same ultrapure water typically used for our intracellular solutions. Finally, samples were concentrated using Vivaspin devices such that sorcin stock solution was ~10 time more concentrated than that used in our experiments.Thus, basal $Ca^{2+}$ levels should not be different from those present in our normal solutions. It is also important to mention that the intracellular solution also included 500 $\mu$M EGTA, which should buffer extra $Ca^{2+}$, even that carried in the final dilution of 3 $\mu$M sorcin.

**Reviewer #3 (General assessment and major comments (Required)):**

**The focus of this manuscript from Moglie et al is to investigate calcium entry in post-hearing OHCs via the activation of either voltage-gated calcium channels or the MOC efferent fibers. Based on the literature reported, very little is known about how OHCs handle increases in cellular calcium, although oncomodulin is believed to be the major calcium buffer in these cells. Therefore, this work attempts to address this gap in our knowledge by using a combination of calcium imaging and electrophysiology. From the results presented, the authors conclude that the large calcium signals generated by the opening of calcium channels appear to be modulated by ryanodine receptors. In addition, the opening of nicotinic receptors, caused by ACh released from active efferent fibers produced calcium transients that were contained by cisternal calcium-ATPases. The authors have also provided results that sorcin, a calcium binding protein involved in controlling calcium in myocytes, appears to control basal calcium levels and MOC synaptic activity in OHCs. The topic of the study is very interesting but unfortunately there are several major shortcomings in the design and execution of the work that drastically lower its impact. Moreover, the work appears to be designed and written for a specialized "auditory" audience.**

We appreciate that the reviewer found this topic interesting, but regret that the impact of the paper could be diminished due to what the reviewer considered ‘major shortcomings’. We hope to address the reviewer’s concerns in this rebuttal.

**The main issue of the paper is that the imaging data is used as the primary means of quantifying calcium changes under different experimental conditions, including the measurements of basal calcium level. However, all experiments were performed with a non-ratiometric calcium dye, making most of the conclusions and assumptions extremely difficult to interpret.**

It is difficult to judge with this sole comment why the author considers that ratiometric dyes should be used to quantify **relative** changes in $Ca^{2+}$ levels. A large proportion of $Ca^{2+}$ imaging studies have been done with non-ratiometric dyes. To name a few that have been critical to the field (and are cited in our study): Beurg et la., 2009; Beurg et al., 2010; Tucker & Fettiplace, 1995; Mammano et al., 1999; Lelli et al., 2003; Frolenkov et al., 2000; Evans et al., 2000). Also, others not cited in our work, such as: Frank et al., 2009 PNAS or Kennedy, 2002 Cell Calcium.
Actually, we intended to use ratiometric dyes when we first tried to image $Ca^{2+}$ transients through $\alpha9\alpha10$. But the signal to noise ratio was very low, and it was difficult to resolve single synaptic events (such as those in Fig. 2, and also Fig. 1 in Moglie et al., 2018) and even trains. These are the main reason we finally decided to use Fluo-4.
Further below we address issues concerning basal calcium measurements.

**Another problem is that the authors make very specific conclusions regarding the mechanisms involved in calcium handling in OHCs, which are used to explain/understand how OHCs operate in vivo. However, experiments were done using whole-cell patch clamp, which is far from physiological, using unphysiological voltages (-100 mV) and at room temperature. The authors should provide evidence that the mechanisms proposed using the above experimental conditions are physiologically relevant.**

These are important points that may require further explanation. We regret that Reviewer #3 has found issues on the interpretations of our experimental results. We will refer to some of the technical flaws that the reviewer pointed out. Firstly, as it was indicated on line 99 in Results (and further clarified in Methods), we only switched the OHCs Vm to -100 mV for a short period of time during efferent stimulation protocol, with the purpose of increasing $Ca^{2+}$ driving force. Otherwise, the cell was voltage clamped at -40 mV, including the resting period between repetitions of MOC stimulations (5-30 sec) and also when checking series resistance, and when stimulation protocols were changed (~ minutes). In other words, over minutes of recordings (4 to 30 minutes), OHCs were clamped at -100 mV for < 5 %. Therefore, considering that resting membrane potential could strongly influence cytoplasmic $Ca^{2+}$ concentration, cistern load, etc.; we believe that our stimulation protocol would only marginally affect $Ca^{2+}$ homeostasis at rest.
As reviewer surely knows, tissue degenerates very quickly at the age we recorded from, and not more than one OHCs recording could be done per preparation. Intense illumination with LED light imposes an additional experimental difficulty due to the release of free radicals within the cell cytoplasm (partially alleviated by the addition of ascorbic acid to the intracellular solution). At body temperature, these problems only worsen and most of the times recordings are simply lost in a few minutes. In the past, however, we have managed to complete a limited data set of OHCs experiments at near body temperature and did not find any significant difference with room temperature experiments in either size of IPSCs, probability of release, or facilitation (Ballestero et al., 2011). Thus, we consider that there is no real need to repeat any of our experiments at body temperature.
On the other hand, we do not see how these experiments could be done without patch-clamp recordings, which was additionally used to deliver drugs intracellularly.

**Figures 1 and 2 describe the same aspect, and should be combined. Also, it is not clear why 1 mM Ach was used for the experiments. How do the authors know that this is a physiologically saturating concentration?**

In our opinion Figure 1 provides a good introduction to the rest of the study, further strengthened by the inclusion of a new panel A with an illustration of the focal plane on an OHCs, as requested by one of the reviewers. In addition, the size of the $Ca^{2+}$ signals due to local ACh applications is critical for the concept of non-saturating $Ca^{2+}$ indicator in the rest of the study (see response to the next point raised by the reviewer). Thus, we would rather keep Figs. 1 and 2 in the current format.
From previous work in OHCs and IHCs (Lioudyno et al., 2004; Gómez-Casati et al., 2005) we know that the $\alpha9\alpha10$ $EC_{50}$ for ACh is ~100 $\mu$M. According to our current experiments, 300 $\mu$M ACh produced a current that was only 4% smaller than that of 1 mM (340 $\pm$ 57 pA and 353 $\pm$ 53 pA, respectively, Vh = -40mV, n = 4 – 6). Therefore, we conclude that 1 mM is nearly saturating. Now it reads in line 79:
*"The application of a nearly saturating concentration of ACh (1 mM) (Gómez-Casati et al., 2005; Lioudyno et al., 2004) produced a global and long lasting elevation of cytoplasmic $Ca^{2+}$"*

**Figure 3G-H highlights another major issue with the method used. The similarity of the calcium change between the different stimulus durations could just be due to dye saturation, which is in fact suggested by the initially flat response in panel G despite the reduction in current. This finding should be corroborated by evidence indicating that the calcium dye is not saturated under their experimental conditions.**

This issue was addressed due to a similar request by reviewer #1. We will transcribe the response here:

We regret that this issue was not sufficiently clarified in our original version of the manuscript. It should firstly indicated that a high affinity indicator was selected because of the small $Ca^{2+}$ influx generated during physiological $\alpha9\alpha10$ activation, that was on the verge of our detection threshold. A lower affinity dye would not be able to resolve those $Ca^{2+}$ transients. This important observation is now clarified on line 95 on Results:

*"$\alpha9\alpha10$ receptors are highly $Ca^{2+}$ permeable, but due to the much higher concentration of $Na^+$ and $K^+$ in the extracellular solution (150 mM together vs 1.3 mM $Ca^{2+}$, see Methods), the majority of the current allowed in physiological conditions is due to monovalent cations (Weisstaub et al., 2002). To circumvent the small $Ca^{2+}$ influx, we adopted two technical measures. Firstly, a high affinity $Ca^{2+}$ indicator, such as Fluo-4, was used to detect synaptic $Ca^{2+}$ transients. And secondly, to maximize the $Ca^{2+}$ driving force, OHCs were transiently voltage-clamped at -100 mV..."*

Reviewer #1 also brought up the issue of saturation of the $Ca^{2+}$ dye. We believe that it is very unlikely that Fluo-4 is saturated in experiments included in Figs. 3-7 considering that $Ca^{2+}$ signals detected with the same indicator (and also repeating all meaningful conditions) but using ACh local application (Fig. 1) were ~20 times larger (compared to 80 Hz trains). The integral of the ACh response was >30 times larger than 80 Hz trains. It is likely that a high affinity indicator such as Fluo-4 is saturated in experiments with large, and sustained, $Ca^{2+}$ influx as those obtained in ACh applications. But the currents are significantly smaller, and briefer, during trains of MOC stimulations, indicating that Fluo-4 is within its dynamic range of $Ca^{2+}$ sensitivity. Moreover, the linear correlation between $\Delta F/F$ and charge in Fig. 3E points in the same direction.
To clarify this problem, a sentence was included in Results, line 144:

*"Comparing $Ca^{2+}$ signals in Figs. 1 and 3 it becomes clear that synaptic activation of $\alpha9\alpha10$ receptors during trains is (at least) ~20 times smaller than that obtained with locally applied ACh. This could be due to the rather low release probability of MOC terminals (Ballestero et al., 2011; Vattino et al., 2020), that our stimulation paradigm typically activates a single axon (see Methods), and that activation by ACh does not rely of vesicle availability. This drastic difference in measurable $\Delta F/F_0$ amplitude between ACh application and electrical stimulation also suggests that the $Ca^{2+}$ indicator used in these experiments, Fluo-4, were not saturated during synaptic trains. The tight correlation between $\Delta F/F$ and charge in Fig. 3E further supports this conclusion."*

And also in the Discussion section, line 272:
*"Synaptic $Ca^{2+}$ signals of both origins were largely smaller than those obtained by local perfusion of a high ACh concentration (compare Figs. 1, 3 and 6), suggesting that the $Ca^{2+}$ indicator used in these experiments (Fluo-4) was not saturated during physiological activation. The selection of the indicator was based upon the signal to noise ratio obtained in experiments that sought to detect $Ca^{2+}$ influx through the highly $Ca^{2+}$ permeable $\alpha9\alpha10$ receptors, which actually allows predominantly $Na^+$ through when activated in physiological*

*conditions (140 mM $Na^+$ and 1.3 mM $Ca^{2+}$, as in our experiments) (Moglie et al., 2018; Weisstaub et al., 2002).”*

**Figure 4 describes an even more problematic result. Here calcium changes are reported as DF instead of DF/F0, which is highly inappropriate as it makes comparing different recordings extremely unreliable (F0 can vary significantly between experiments, see Figure 4F). Similarly, DF measurements are done in other experiments (e.g. Figure 5D), in which data for the control condition comes from a different cell. As mentioned above, this problem could be avoided by using a ratiometric dye (e.g., fura-2 or furaptra see Beutner-Moser 2000?).**

$\Delta$F, instead of $\Delta$F/F0, was only used in the case of experiments in which manipulations with pharmacological agents produced a long lasting change in basal $Ca^{2+}$. Reporting un-normalized fluorescence signals is not rare and has been done in multiple studies, particularly in the cochlea (Kennedy, 2002; Frank et al., 2009; Moglie et al., 2018). It is also important to highlight that all these cited studies used non-ratiometric dyes. The study by Kennedy (2002) is of particular importance because a similar approach with reporting $\Delta$F was applied in experiments using thapsigargin (and other ATPases blockers), which produced an increase in basal $Ca^{2+}$.

As the reviewer may know, dividing changes in fluorescence ($\Delta$F) by the basal values ($F_0$) helps in normalizing cells with different indicator load, or regions of a given cell where dye diffusion is not homogeneous. None of these situations are found in our experimental conditions, given that the dye was always introduced into the cell at the same concentration by the patch pipette, and also because OHCs are small and their basolateral compartment is not diffusionally restricted.

The limitations in using ratiometric dyes to image the small $\alpha 9\alpha 10$ mediated $Ca^{2+}$ events have been mentioned above.

**Figure 5B. It is surprising to see that a similar variance in baseline calcium level to that reported in Figure 4E (again using non-ratiometric measurements), is now just significant and used to support one of the main conclusions of the paper. Considering that the method used does not provide quantifiable baseline calcium levels, how are the authors able to exclude bias in their measurements due to experimental variability? What is the biological replica needed to validate their statistics based on the mean +/- sem? Also, the fact that adding sorcin "increases" the resting calcium level does not prove that it has a role in OHC function; it only shows that sorcin affects calcium levels, which is not surprising since it is a calcium binding protein.**

We appreciate reviewer’s effort in looking at our dataset in detail, which gives us the opportunity to better explain the results. Statistical analysis was not clearly explained in our previous version of the manuscript, but we now included a whole section in Methods.

Statistics was performed using the most conservative approach given the sample size, i.e. non-parametric methods. Moreover, the effect size analysis was included on our first submission so reviewers could judge differences independently of the statistical significance.

Regarding variability and datasets in Figs. 5B and 4F it should be considered that datasets were obtained in different cells due to the nature of the experiment, but with highly similar recording conditions (all conditions were identical but the addition of sorcin, ryanodine, dantrolene, or DMSO as vehicle). But this fact neither represented a bias by itself, nor prevented us from making a statistical comparison (see line 187 for ryanodine, and 199 for sorcin). In the case of changes in basal $Ca^{2+}$ due to sorcin (Fig. 5), we further evaluated leak current values in each dataset. As the reviewer may know, leak current can reveal bad recording conditions, which in turn could produce an increase in basal $Ca^{2+}$ concentration. To rule out a bias in sorcin values due to bad (or worse) recording conditions, we further analyzed $Ca^{2+}$ levels as a function of leak current for each cell. As indicated in line 203 of Results, we found that sorcin produced an increase in resting $Ca^{2+}$ levels irrespective of recording conditions.

The effect of sorcin on the resting $Ca^{2+}$ level was actually not expected. Sorcin is a $Ca^{2+}$ binding protein, but its proven role in cardiomyocytes is through the modulation of RyR, channels and pumps. Not as a buffer *per se*. If it only operated as a buffer, the addition of sorcin to the OHCs cytoplasm would result in a decrease, not a rise in resting $Ca^{2+}$ concentration. And most importantly, the concentration of sorcin added to the intracellular solution could only minimally contribute to a $Ca^{2+}$ buffering effect, since it only represents a 0.3% of the total $Ca^{2+}$ buffer used in our intracellular solution. All these facts are now discussed in the new version of the manuscript, in line 376:

"*The concentration of sorcin used in our experiments matches those in studies showing effects on RyR, $Ca^{2+}$ channels, and Na+-$Ca^{2+}$ exchangers (1-5 $\mu$M) (Farrell et al., 2003; Fowler et al., 2009; Lokuta et al., 1997; Zamparelli et al., 2010). Independently of its mechanism of action, our experiments indicate that sorcin is not merely a $Ca^{2+}$ buffer. According to our experiments, the amount of additional buffer that sorcin represent in the electrode solution is negligible, < 0.5% (3 $\mu$M sorcin in a solution with 400 $\mu$M Fluo-4 and 500 $\mu$M EGTA), precluding a mere buffering effect. Moreover, the reported concentration of oncomodulin and other proteinaceous buffers in OHCs cytoplasm (in the mM range) (Hackney et al., 2005) would largely out compete the buffering capacity of a few $\mu$M concentration of sorcin.*"

**Figure 7D is a bit puzzling to me but I may have missed some underlying reason from published work. Why do Ryn concentrations that are known to either facilitate or block the receptors cause the same change in calcium level?**

The only statistically significant difference in Figure 7D is between control and 1 $\mu$M ryanodine. The rest does not reach statistical significance. $Ca^{2+}$ transients recorded by step depolarizations in 100 $\mu$M ryanodine were very similar to control, with the exception of one cell showing larger size (see individual symbols in Fig. 7D).

**The method section should contain a statistical statement. It should also explain the reason for using non-parametric analysis for the statistical comparisons. Also, most of the methods are only briefly described; although the authors have probably published these methods before, the method section should be more self-explanatory e.g. exactly how was the photobleaching correction performed?**

We acknowledge that our Methods section lacked some important information such as the justification of the statistical tests used in the study. The new version of Methods is more detailed, and includes this and other important facts. In particular, photobleaching was corrected by fitting a line pre-stimulus baseline and the final fluorescence (line 465, Methods).
